# Repurposing polyamines to prevent arrhythmias in Short QT Syndrome type 3, a potentially lethal disease

**DOI:** 10.1101/2025.09.10.675300

**Authors:** Ana I. Moreno-Manuel, Francisco M. Cruz, Álvaro Macías, Eva Cabrera-Borrego, José Jalife

## Abstract

Short QT Syndrome type 3 (SQTS3) is a channelopathy characterized by the abbreviation of the QT interval on electrocardiogram and life-threatening arrhythmias. SQTS3 is caused by gain-of-function mutations in *KCNJ2*, which codes the strong inward rectifier potassium channel Kir2.1 responsible for the repolarizing current I_K1_. Inward-going rectification of I_K1_ is due to a voltage-dependent block by intracellular magnesium and polyamines. We evaluated the therapeutic potential of extrinsic polyamine administration to restore normal cardiac repolarization and prevent life-threatening arrhythmias in an AAV9-mediated mouse model of SQTS3 carrying the Kir2.1^M301K^ mutation, which shortens the QT interval to a minimum of 194ms in patients. Compared to Kir2.1^WT^, the QT interval was significantly shortened in Kir2.1^M301K^ mice. Intracardiac stimulation induced long-lasting, high-frequency ventricular tachycardia. Patch-clamping demonstrated an extremely abbreviated action potential duration (APD) in Kir2.1^M301K^ cardiomyocytes due to a Kir2.1 lack of inward rectification. These I_K1_ defects decreased I_Na_ density and Na_V_1.5 expression. The mechanistically targeted Kir2.1 blockage by exogenous polyamines (spermine intraperitoneally or spermidine orally) prolonged the APD and the QT interval, and significantly reduced arrhythmia inducibility in Kir2.1^M301K^ mice. Polyamines significantly reduced the I_K1_ gain-of-function, an effect that alleviated the I_Na_ reduction in mutant cardiomyocytes. Therefore, repurposing polyamine administration might be a novel and effective therapeutic strategy for managing cardiac arrhythmias in SQTS3 patients.

## INTRODUCTION

Short QT Syndrome (SQTS) is a rare, but highly lethal inheritable channelopathy characterized by an abnormally short QT interval on the electrocardiogram (ECG) and an increased risk for atrial and life-threatening ventricular arrhythmia (AF, VT/VF). SQTS is one of the frequent syndromes associated with sudden cardiac death (SCD),^1^ which is often the first manifestation of the disease with devastating consequences to the patient and their family. Approximately 250 cases have been diagnosed in nearly 150 families worldwide, all during the last two decades.^2–4^ Despite their heterogeneous phenotype, the penetrance of the SQTS causative mutations is remarkable and patients experience a poor quality of life.^5^ Therefore, there is an urgent need to improve the diagnosis and prevent SCD of SQTS patients.

The molecular mechanisms leading to channel dysfunction, cardiac rhythm disturbances, and related disorders associated with SQTS remain incompletely understood. SQTS-linked variants in *KCNH2* (SQTS1), *KCNQ1* (SQTS2), *KCNJ2* (SQTS3), and *SLC4A3* (SQTS8) are the only validated genotype-positive SQTS subtypes to date.^6, 7^ The SQTS1-causative mutations hERG^T618I^ and hERG^N588K^ are the most common (25.9% of genotyped probands) and second most common (18.5%), respectively, of the clinically occurring SQTS variants.^8, 9^ However, the SQTS3-leading mutation Kir2.1^M301K^ causes the most pathogenic QT abbreviation.^10, 11^ Kir2.1^M301K^ was described in an 8-year-old girl with a QTc interval of 194ms, paroxysmal AF and VF inducibility, also suffering from multiple disorders, such as severe mental retardation, abnormal proliferation of esophageal blood vessels, epilepsy, and Kawasaki disease.^11^ This patient suddenly died ten years after diagnosis. The methionine 301 in Kir2.1 forms a pore-facing loop region,^12^ and functional analysis in heterologous expression systems revealed that its replacement with a lysine abolished I_K1_ in homozygosis and increased outward I_K1_ in heterozygosis.^11^

SQTS3 is caused by mutations in the *KCNJ2* gene that codes Kir2.1, the strong inward rectifier potassium channel. Kir2.1 is responsible for I_K1_, which is essential in maintaining the resting membrane potential (RMP) and the final phase of action potential (AP) repolarization.^13–15^ Kir2.1 conductance changes (rectifies), and the slope of its current/voltage (IV) relationship is modified at voltages positive to the K^+^ equilibrium potential, being the current abolished at voltages close to 0mV. Inward rectification of Kir2.x channels results from blockade by intracellular magnesium and polyamines (spermine, spermidine and putrescine),^16–18^ which upon membrane depolarization penetrate the cytoplasmic pore of the channel, binding to specific negatively charged residues and obstructing the outward flow of potassium.^19–21^ Spermine is the main channel blocker responsible for inward rectification, followed by spermidine, putrescine, and magnesium.^16, 18, 22–24^ Kir2.1 channels possess multiple polyamine binding sites.^25^ They are negatively charged regions, one in the transmembrane domain, involving D172, and the other in the cytoplasmic region, involving E224, D255, D259 and E299.^12^ The initial step in the rectification process involves the interaction of polyamines with E224 and E299 located in the COOH-terminus of the Kir2.1 channel.^18, 22, 26^ The second step is more voltage-dependent and implies the binding of polyamines at the D172 residue,^22,26^ which is deeper inside the Kir2.1 pore. Mutations at any of the above residues result in a defective inward rectification of I_K1_. Polyamines are non-toxic and have been shown to be important in ageing, cancer and other diseases, but inward rectification at depolarized potentials is likely their most important function.^27^ However, to our knowledge, polyamines have never been considered as a therapeutic option in SQTS3.

The development of innovative mechanistically targeted therapies for individuals suffering from SQTS3 is urgent. Currently, treatment for SQTS patients is ill-defined and there are no complex, standardized or comprehensive clinical protocols of action guiding the patients’ management and the arrhythmias prevention. The rarity of the disease and its recent description have prevented the design of clinical trials on appropriate groups of patients.^4^ In symptomatic individuals in whom an implanted cardioverter defibrillator (ICD) is not recommended, pharmacological treatment is mandatory. However, the currently available drugs have loads of out-off-target effects because they interfere with the behaviour of various ion channels and their regulators. Recommended drugs are antiarrhythmics like ibutilide, flecainide, sotalol, amiodarone, propafenone, and β- blockers, like metoprolol and carvedilol.^4^ β-blockers reduce adrenergic input to the heart, a mechanism that helps control the heartbeat and prevent symptoms.^28^ The latest European Society of Cardiology (ESC) Guidelines for the management of patients with ventricular arrhythmias and prevention of SCD recommend an ICD and treatment with quinidine.^29^ It is known that quinidine blocks I_Na_, decreasing phase 0 of the action potential (AP); it also reduces I_Kr_, I_Ks_, I_K1_, I_to_, I_Ca-L_ and I_NaLate_, leading to AP duration (APD) prolongation.^30^ Both quinidine and hydroxyquinidine are tested as pharmacological treatments of SQTS patients because they prolong the QTc interval, reducing the incidence of life-threatening arrhythmias.^31–33^ Nevertheless, in some countries, quinidine has been removed from the market because of important side-effects. On this basis, the design of new, safer and more effective drugs is urgent to improve life-quality and prevent arrhythmias and SCD in SQTS patients.

This original article documents our experimental results on the underlying mechanisms of SQTS3 caused by Kir2.1^M301K^, the most pathogenic SQTS variant described in the literature.^11^ We report on data derived from the first *in-vivo* adeno-associated (AAV)- based mouse models of cardiac-specific SQTS3 mimicking the electrophysiological phenotype of patients expressing Kir2.1^M301K^. The availability of cardiac-specific SQTS3 mouse models helped identify new molecular targets for the design of novel antiarrhythmic drugs in an inheritable cardiac disease that currently has no defined therapy. Our results demonstrate that polyamines effectively correct the APD, QT interval, and reduce arrhythmogenesis in SQTS3 mice, and suggest that repurposing these naturally occurring molecules might be a new therapeutic option to prevent life-threatening arrhythmias in SQTS3 patients.

## MATERIALS AND METHODS

Detailed descriptions are provided in the **Supplementary Materials**.

### Study Approval

All experimental procedures using animals conformed to EU Directive 2010/63EU and Recommendation 2007/526/EC, enforced in Spanish law under *Real Decreto 53/2013*. The local ethics committees and the Animal Protection Area of the Comunidad Autónoma de Madrid (PROEX 226.5/23) approved animal protocols.

### Mice

Four-week-old C57BL/6J male mice were obtained from Charles River Laboratories. Mice were reared and housed following institutional guidelines and regulations.

### Adeno-associated virus (AAV) production, injection, and mouse model generation

Vectors driven by the cardiomyocyte-specific cardiac Troponin-T proximal promoter (cTnT), and encoding wildtype Kir2.1 (Kir2.1^WT^) or the SQTS3 Kir2.1^M301K^ mutant were packaged into AAV serotype 9 (AVV9) capsids.^34–37^ After anesthesia (Ketamine 60 mg/kg and Xylazine 20 mg/kg i.p.), 4- to 5-week-old mice were administered 3.5x10^10^ viral genomes (vg) per animal i.v.^34, 38^ Mice were used for experiments 8 weeks after the AAV9 infection.

### Polyamine administration

Fifteen- to 17-week-old male mice were used to test the effects of polyamine administration. Ten mg/Kg spermine was administered i.p. daily for 10 or 20 days, changing the administration area to avoid inflammation.^39^ Three mM spermidine was administered via drinking water for 10 or 20 days, or a maximum of 6 months, changing the water and preparing fresh spermidine every 3 days.^40^ Fifty-two-week-old mice also underwent long-term treatment with spermidine.

### Surface ECG recordings

Mice were anesthetized with 0.8-1% Isoflurane in oxygen. A subcutaneous 23-gauge needle electrode connected to an MP36R amplifier (BIOPAC Systems) was attached to each limb, and six-lead surface ECGs were recorded. We analyzed blindly the recordings using AcqKnowledge 4.1 and LabChart softwares.

### *In-vivo* intracardiac electrophysiology

After anesthesia (Ketamine 60 mg/kg and Xylazine 20 mg/kg i.p.), an octopolar catheter (Science) was inserted in the heart through the jugular vein, as described previously.^41, 42^ Ventricular refractory periods and arrhythmia inducibility were assessed in control and mutant mice.

### Ventricular cardiomyocyte isolation

After cervical dislocation, the mouse heart was cannulated through the ascending aorta, mounted on a Langendorff-perfusion apparatus and retrogradely perfused as per Macías *et al*.^43^

### Patch-clamping of ventricular cardiomyocytes

Whole-cell (current and voltage clamp) patch-clamping and data analysis procedures were described previously^44–48^ (see **Supplementary Materials**). External and internal solutions are listed in **Supplementary Table 1**.^47, 49^

### Statistical analysis

Data are expressed as mean ± SEM, and differences are considered significant at p<0.05 (*p<0.05; **p<0.01; ***p<0.001; ****p<0.0001). Note that “N” refers to the number of mice, and “n” to the number of cells analyzed per mouse.

## RESULTS

### Design and generation of the mouse SQTS3 model

We generated a mouse model of SQTS3 using specific constructions containing the mutated versions of *KCNJ2* (Kir2.1^M301K^) packaged in AAV9 (**Supplementary Figure 1Ai**). AAV9-generated Kir2.1^WT^ animals were used as controls. Ex-vivo hearts analysis confirmed the cardiac expression of Kir2.1^WT^ and Kir2.1^M301K^ through fluorescence emitted by the tdTomato reporter present in the AAV constructs, which was totally absent in untransfected hearts (**Supplementary Figure 1Aii**). Haematoxylin-Eosin staining revealed that WT and mutant mouse hearts were structurally normal. Immunohistochemistry and immunohistofluorescence with tdTomato in cardiac slices confirmed a mosaic and wide AAV9 distribution throughout myocardium (**Supplementary Figure 1iii-iv**), as reported previously.^34^ Total Kir2.1 protein levels in the hearts of mice expressing Kir2.1^WT^ and Kir2.1^M301K^ were very similar to untransfected hearts (**Supplementary Figure 1B**, left). We also confirmed that there was no increase in Kir2.1 expression at the plasma membrane of Kir2.1^M301K^ compared to Kir2.1^WT^ cardiomyocytes (**Supplementary Figure 1B**, right).

Echocardiographic analysis confirmed normal structure and contractile function of both Kir2.1^WT^ and Kir2.1^M301K^ hearts (**Supplementary Figure 2**). We previously reported that the echocardiographic values of untransfected animals were similar to AAV9-Kir2.1^WT^ control mice,^50^ and that genetic haploinsufficiency did not operate after expression in *trans* of Kir2.1^WT^ and Kir2.1^SQTS3^.^51^ We also checked that ECG, arrhythmia inducibility and cellular electrophysiological parameters (e.g., APD, I_K1_ and I_Na_) of Kir2.1^WT^ were similar to untransfected mice (see **Supplementary Figure 3**). Therefore, since AAV9- Kir2.1^WT^ did not alter total Kir2.1 expression and represented a transfected control that did not disturb the murine electrical phenotype, all subsequent experiments were conducted in Kir2.1^M301K^ animals and compared with Kir2.1^WT^ controls.

### The QT interval is significantly abbreviated in Kir2.1^M301K^ mice

We recorded ECGs from Kir2.1^WT^ and Kir2.1^M301K^ mice. Whether non-corrected or corrected by the Bazett or the Framingham formula,^52, 53^ mean QT intervals in SQTS3 were all shorter than Kir2.1^WT^ mice (Kir2.1^WT^, 30.1±1.2 ms > Kir2.1^M301K^, 16.4±0.7 ms) (**Figure 1A-C**). Therefore, the Kir2.1^M301K^ mouse recapitulated the SQTS3 phenotype in that both patient and mouse ECGs demonstrated extreme QT interval shortening.^11^

**Figure 1.**
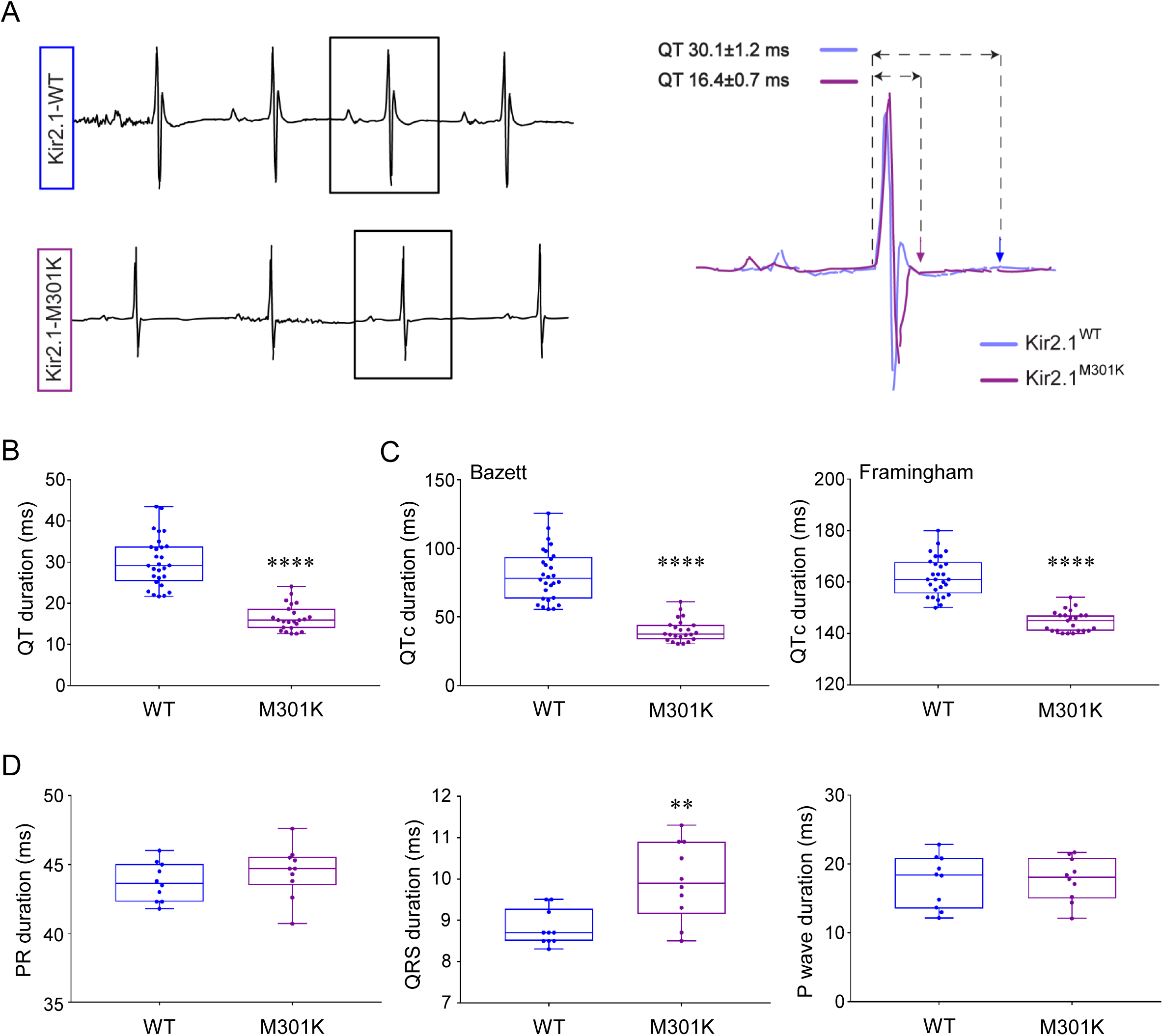
QT intervals in Kir2.1^M301K^ mouse models are significantly abbreviated and the QRS complex prolonged compared to Kir2.1^WT^ mice. A, Representative ECG traces showing QT interval abbreviation in Kir2.1^M301K^ models (purple) compared to Kir2.1^WT^ mice (blue). B, Distribution of QT interval values in all models analyzed (Kir2.1^WT^: 30.1±1.2ms, N=28; Kir2.1^M301K^: 16.4±0.7ms, N=23). C, <u>Left</u>, QTc intervals of Kir2.1^WT^ and Kir2.1^M301K^ mice (80.3±3.5ms and 40.2±1.7ms, respectively; N=23-28) using the Bazett’s equation. <u>Right</u>, QTc intervals in Kir2.1^WT^ and Kir2.1^M301K^ animals (162.2±1.4ms and 144.7±0.8ms, respectively; N=23-28) applying the Framingham’s correction. D, from left to right, PR interval, QRS complex, and P wave duration in Kir2.1^WT^ and Kir2.1^M301K^ mice are shown (N=10 animals per condition). Mann-Whitney test (QT/QTc intervals, PR intervals and QRS complexes) and unpaired 2-tailed Students’t-test (P waves) were used.

As illustrated in **Figure 1D**, the PR interval and P wave duration were unaffected by the mutation. However, the QRS was prolonged in the ECG of Kir2.1^M301K^ mice, indicating that intraventricular conduction was slower than Kir2.1^WT^ mice. The significance of such a defect will become apparent below.

### Long-lasting and high-frequency ventricular tachycardia in Kir2.1^M301K^ mice

We used an intracavitary octopolar catheter and intracardiac programmed electrical stimulation protocols to characterize the *in-vivo* cardiac electrophysiological properties and arrhythmogenicity of these mice. As expected, the ventricular refractory period was reduced in Kir2.1^M301K^ compared to Kir2.1^WT^ mice (**Supplementary Figure 4**), which was consistent with their different QT intervals.

To determine atrial and ventricular arrhythmia inducibility in each mouse group, we applied S1-S2 trains of 10 and 25 Hz (see **Materials and Methods**) on either the right ventricle (RV) or right atrium (RA) through the octopolar catheter. RV stimulation induced VT/VF in Kir2.1^M301K^ mice (**Figure 2Ai**). Seven out of 8 Kir2.1^M301K^ mice (87.5%) demonstrated the highest percentage of longer-lasting (>5s) and high-frequency VT/VF episodes (**Figure 2B**).

**Figure 2.**
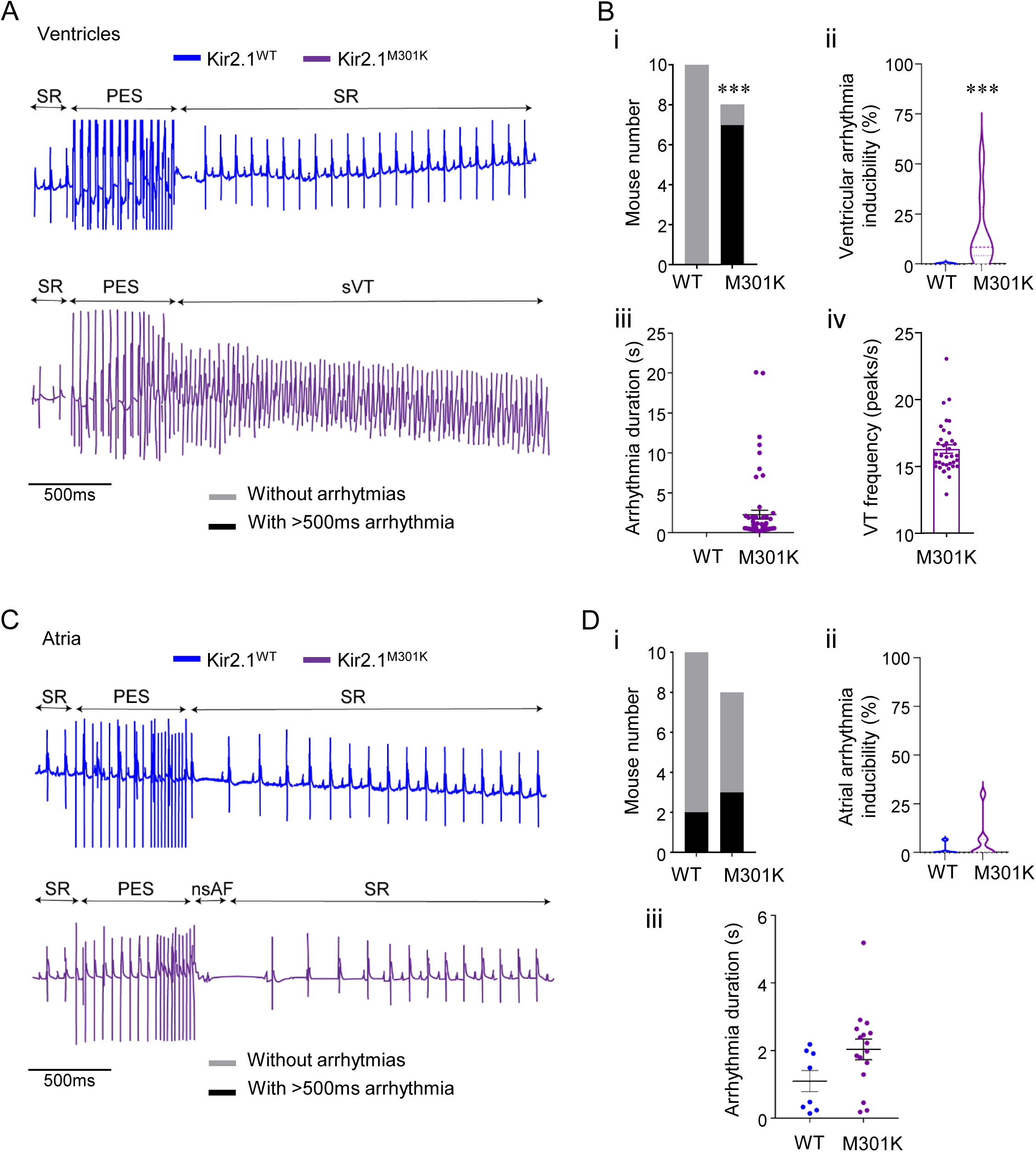
Ventricular arrhythmia inducibility is significantly increased in SQTS3 Kir2.1^M301K^ models. **A, C**, ECG recordings during and after the programmed electrical stimulation (PES) are shown for Kir2.1^WT^ (blue) and Kir2.1^M301K^ (purple) animals. Sinus rhythm (SR), PES, and sustained ventricular tachycardia or non-sustained atrial fibrillation (sVT or nsAF, respectively) after ventricular (**A**) or atrial (**C**) stimulation. **B, D**, **i**, Number of mice inducible for >500ms ventricular (**B**) and atrial (**D**) arrhythmias. **ii**, Violin plots showing the distribution for ventricular (**B**) and atrial (**D)** inducibility presented as the percentage of stimuli that triggered arrhythmia in each group. **iii**, Duration (s) of all ventricular (**B**) and atrial (**D)** arrhythmias recorded in each group while stimulation. **B, iv**, Frequency of the VT episodes lasting more than 500ms recorded in Kir2.1^M301K^ mice measured as peaks per second. Fisher’s exact test (presence/absence of arrhythmias) and Mann-Whitney test were applied (***p<0.001).

On the other hand, focusing on the atria, some Kir2.1^M301K^ animals were AF inducible although the duration of the AF episodes was brief (**Figure 2Aii** and **2C**).

Altogether, these data demonstrated that the mutation Kir2.1^M301K^ causes a pathogenic ventricular arrhythmia phenotype mediated by mechanisms that were consequently studied.

### APD is extremely short in Kir2.1^M301K^ cardiomyocytes

To characterize the electrophysiological properties of ventricular isolated cardiomyocytes, we carried out patch-clamp experiments in the current-clamp mode and recorded APs in both Kir2.1^WT^ and Kir2.1^M301K^ conditions. We demonstrated that the Kir2.1^M301K^ mutation consistently and significantly abbreviated the ventricular APD at almost all repolarization percentages and frequencies studied (**Figure 3A-B** and **Supplementary Figure 5**). Moreover, APD shortening was not rate dependent. However, RMP, *d*V/*d*t and AP amplitude (APA) were not affected by the Kir2.1^M301K^ mutation (**Figure 3C-E**).

**Figure 3.**
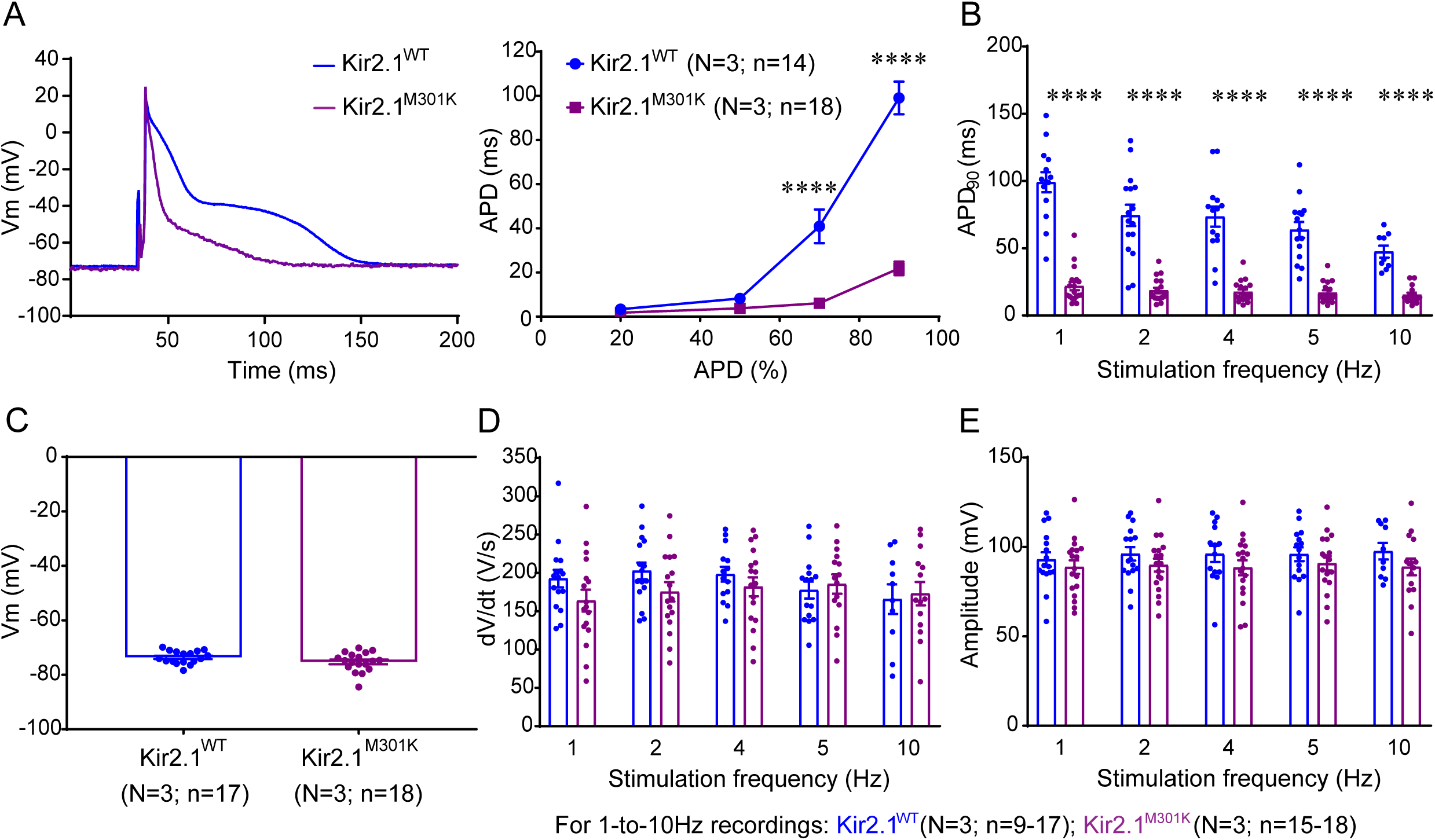
The SQTS3 Kir2.1^M301K^ mutation extremely reduces ventricular APDs. **A**, Representative APs from isolated Kir2.1^WT^ (blue) and Kir2.1^M301K^ (purple) ventricular cardiomyocytes recorded at 1Hz (<u>left</u>), and APDs at 20, 50, 70 and 90 % repolarization (<u>right</u>). **B**, APD_90_ at 1-to-10 Hz are shown for each group following the same color pattern. **C**, RMP (in mV), **D**, maximum upstroke velocity (*d*V/*d*t; V/s), and **E**, AP amplitude (mV) for 1-to-10 Hz stimulation frequencies recorded from Kir2.1^WT^ (N=3, n=9-17) and Kir2.1^M301K^ (N=3, n=15-18) ventricular cardiomyocytes. Unpaired 2-tailed Student’s t-test (RMP), two-way ANOVA (panel A, right), and Mann-Whitney test (for the remaining comparisons) were applied (****p<0.0001).

### I_K1_ is increased and lacks inward rectification, while I_Na_ density and kinetics are altered in Kir2.1^M301K^ cardiomyocytes

We performed whole-cell voltage-clamp experiments to analyze the barium-sensitive I_K1_ in Kir2.1^WT^ and Kir2.1^M301K^ cardiomyocytes (**Supplementary Figure 6**). In Kir2.1^M301K^ cardiomyocytes the inward component of I_K1_ was reduced at voltages negative to -80mV, which decreased the channel slope conductance. But there was also a gain-of-function in the outward component of I_K1_ due to a lack of rectification (**Figure 4A**). Here the channel remained opened at voltages at which it should have been closed (i.e., between -35 and 10 mV).

**Figure 4.**
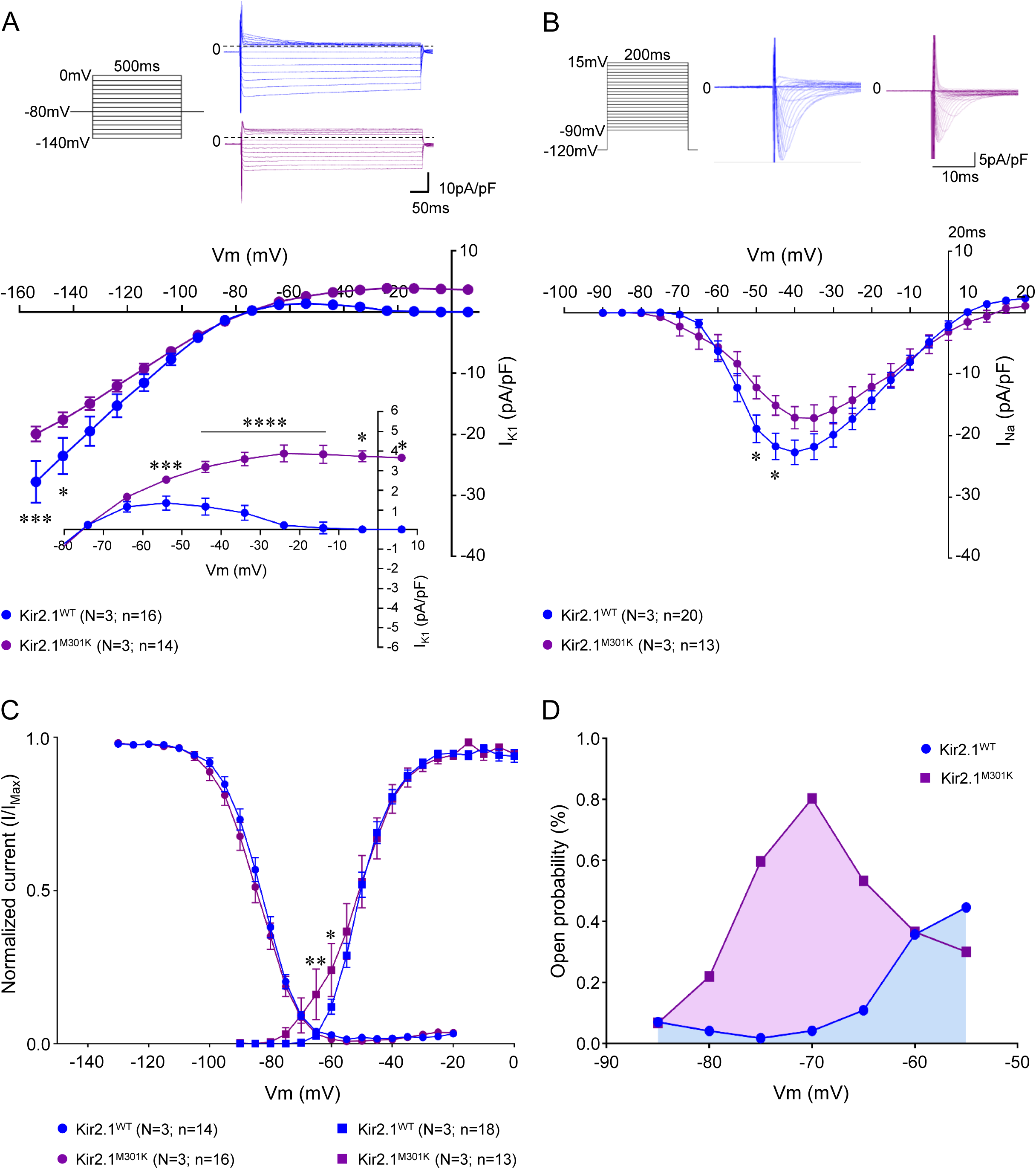
Kir2.1^M301K^ impairs inward-going rectification of I_K1_ and reduces I_Na_ in ventricular cardiomyocytes. **A**, I_K1_ IV relationships for Kir2.1^WT^ (blue; N=3, n=16) and Kir2.1^M301K^ (purple; N=3; n=14) ventricular myocytes. Note the lack of inward-going rectification with increased outward I_K1_ at voltages positive to -55mV (3.83±0.47pA/pF in M301K *vs* -0.12±0.17pA/pF in WT, at -14mV) and loss of inward current at voltages negative to -120mV. **B**, Superimposed I_Na_ IV relationships for Kir2.1^WT^ (N=3, n=20) and Kir2.1^M301K^ (N=3, n=13) isolated cardiomyocytes. The protocols and representative I_K1_ and I_Na_ IV traces for Kir2.1^WT^ (blue) and Kir2.1^M301K^ (purple) are presented. **C**, Na_V_1.5 channel steady-state activation and inactivation curves for Kir2.1^WT^ (N=3, n=14-18) and Kir2.1^M301K^ (N=3, n=13-16). Note the higher open probability at negative voltages in the I_Na_ activation curves of Kir2.1^M301K^ cardiomyocyte. **D**, Na_V_1.5 open probability quantified at window current voltages (from -85 to -55 mV). Two-way ANOVA tests were applied for comparisons (****p<0.0001, ***p<0.001, and *p<0.05).

We wondered whether the arrhythmogenic consequences and the QRS complex prolongation in mice carrying the mutation Kir2.1^M301K^ were due to alterations in Kir2.1 alone or if other channel disturbances were involved. It has been previously described that Kir2.1 form *channelosomes* with the cardiac sodium channel Na_V_1.5 at the plasma membrane, where both channels are reciprocally regulated.^45–47, 50, 54, 55^ Therefore, we performed whole-cell voltage-clamp experiments and, unexpectedly, saw that Kir2.1^M301K^ cardiomyocytes had a reduced I_Na_ density (**Figure 4B**). In addition, while I_Na_ inactivation was unaffected by the mutation, the slope of the I_Na_ activation curve was modified at hyperpolarized potentials causing an increase in the window current (**Figure 4C** and **Supplementary Figure 7**). Both peak I_Na_ density reduction and window current increase are potentially arrhythmogenic,^56^ predisposing the heart to higher VT/VF inducibility in the Kir2.1^M301K^ mice and spontaneous arrhythmias in the SQTS3 patient.

### Na_V_1.5 expression levels are reduced in Kir2.1^M301K^ cardiomyocytes

We aimed to decipher whether the decrease in I_Na_ density was due to a reduced Na_V_1.5 expression. Therefore, we quantified Na_V_1.5 protein in total and plasma membrane-enriched fractions of Kir2.1^WT^ and Kir2.1^M301K^ cardiomyocytes. We observed that the total levels of Na_V_1.5 in hearts expressing Kir2.1^M301K^ were lower than in Kir2.1^WT^ hearts (**Figure 5A**). On the other hand, compared to Kir2.1^WT^, Na_V_1.5 expression was reduced in the plasma membrane-enriched fraction of Kir2.1^M301K^ cardiomyocytes (**Figure 5B**).

**Figure 5.**
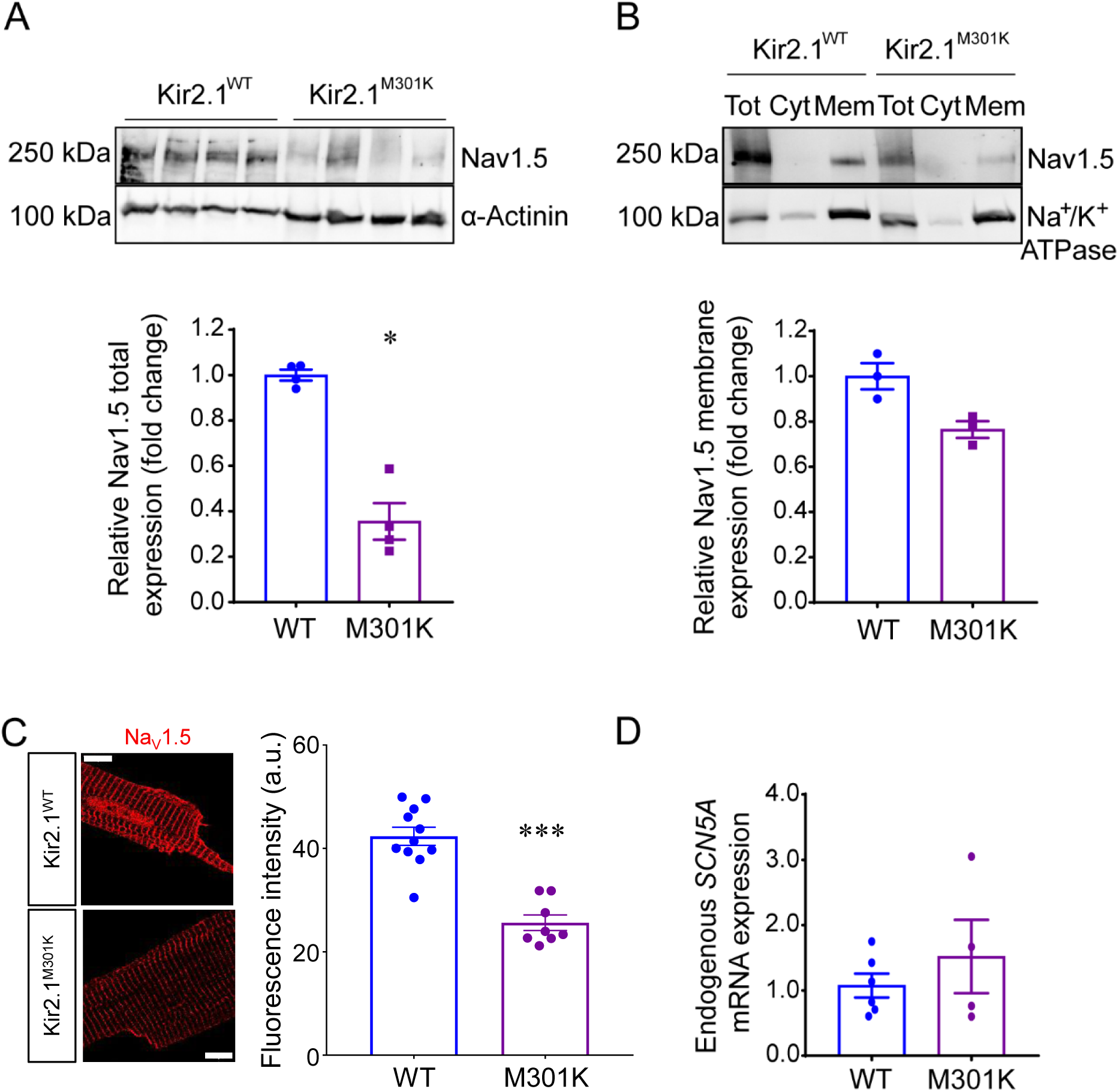
Na_V_1.5 protein expression is reduced in Kir2.1^M301K^ ventricular cardiomyocytes. **A**, Western blot comparing total Na_V_1.5 protein expression in ventricles from Kir2.1^WT^ (blue; N=4) and Kir2.1^M301K^ (purple; N=4) mice. Graph shows western blot quantification normalized using α-Actinin. **B**, Western blot analysis comparing Na_V_1.5 channels in total (Tot), cytosolic (Cyt) and sarcolemmal-enriched (Mem) fractions. The Na^+^/K^+^ ATPase was used as plasma membrane loading control. Graph show western blot quantification of sarcolemmal Na_V_1.5 protein expression in Kir2.1^WT^ and Kir2.1^M301K^ cardiomyocytes (N=3 animals per condition). **C**, Representative confocal images showing Na_V_1.5 channel distribution and fluorescence intensity in Kir2.1^WT^ (N=2; n=11) and Kir2.1^M301K^ (N=2; n=8) expanded ventricular cardiomyocytes. Scale bars, 50μm. Graph shows quantification of Na_V_1.5 fluorescence intensity in both groups. **E**, Quantification of endogenous murine *SCN5A* mRNA expression levels in WT and M301K hearts. Data were normalized by *GAPDH* (N=4-6 animals per condition; experiments performed in triplicate). Mann-Whitney tests were applied for statistics (***p<0.001 and *p<0.05).

Moreover, in expansion microscopy experiments the Na_V_1.5 fluorescence intensity in expanded Kir2.1^M301K^ ventricular cardiomyocytes was lower than Kir2.1^WT^ cells, but its tubular distribution pattern was maintained (**Figure 5C**). However, Na_V_1.5 mRNA expression was similar in Kir2.1^WT^ and Kir2.1^M301K^ cardiomyocytes (**Figure 5D**), indicating the absence of alterations at the transcription level. Thus, the Kir2.1^M301K^ mutation seems to cause alterations in the Na_V_1.5 translation rate.

Apart from Na_V_1.5, we studied the distribution of the already known Kir2.1-Na_V_1.5 *channelosome* interactors SAP97 and α1-syntrophin, and revealed they were similar in Kir2.1^WT^ and Kir2.1^M301K^ cardiomyocytes (**Supplementary Figure 8**).

### Exogenous spermine prolongs the QT interval in Kir2.1^M301K^ mice

After studying the mechanism underlying the I_K1_ increase caused by Kir2.1^M301K^, we hypothesized that polyamines administered systemically could therapeutically restore the QT interval by inducing inward-going rectification of Kir2.1^M301K^ mutant channels. To prove this concept, we treated Kir2.1^M301K^ mice with spermine, the most potent voltage dependent Kir2.1 blocker.^16, 18, 22–24^ Fifteen- to 17-week-old Kir2.1^WT^ and Kir2.1^M301K^ mice were administered 10mg/Kg/day spermine (i.p.) for 10 and 20 days. As previously reported,^39^ after 10 days of treatment, all animals, independently of the genotype, manifested a loss of about 7.5% of their body weight and then stabilized throughout the 20-day duration of the treatment (**Supplementary Figure 9**). Most importantly, the QT/QTc interval was significantly prolonged in mutant mice, without alterations in other ECG parameters (**Figure 6A** and **Supplementary Figure 10A**). Neither the Kir2.1^M301K^ mice nor the Kir2.1^WT^ mouse were inducible for long-lasting (>1s) atrial or ventricular arrhythmias after 10-day spermine treatment (**Figure 6B-D**), suggesting a potential therapeutic action of the drug for the prevention of Kir2.1^M301K^-associated arrhythmogenic events. Surprisingly, only 1 out of 8 Kir2.1^M301K^ mice presented a couple of long-lasting atrial episodes after 20-day treatment (**Figure 6C**).

**Figure 6.**
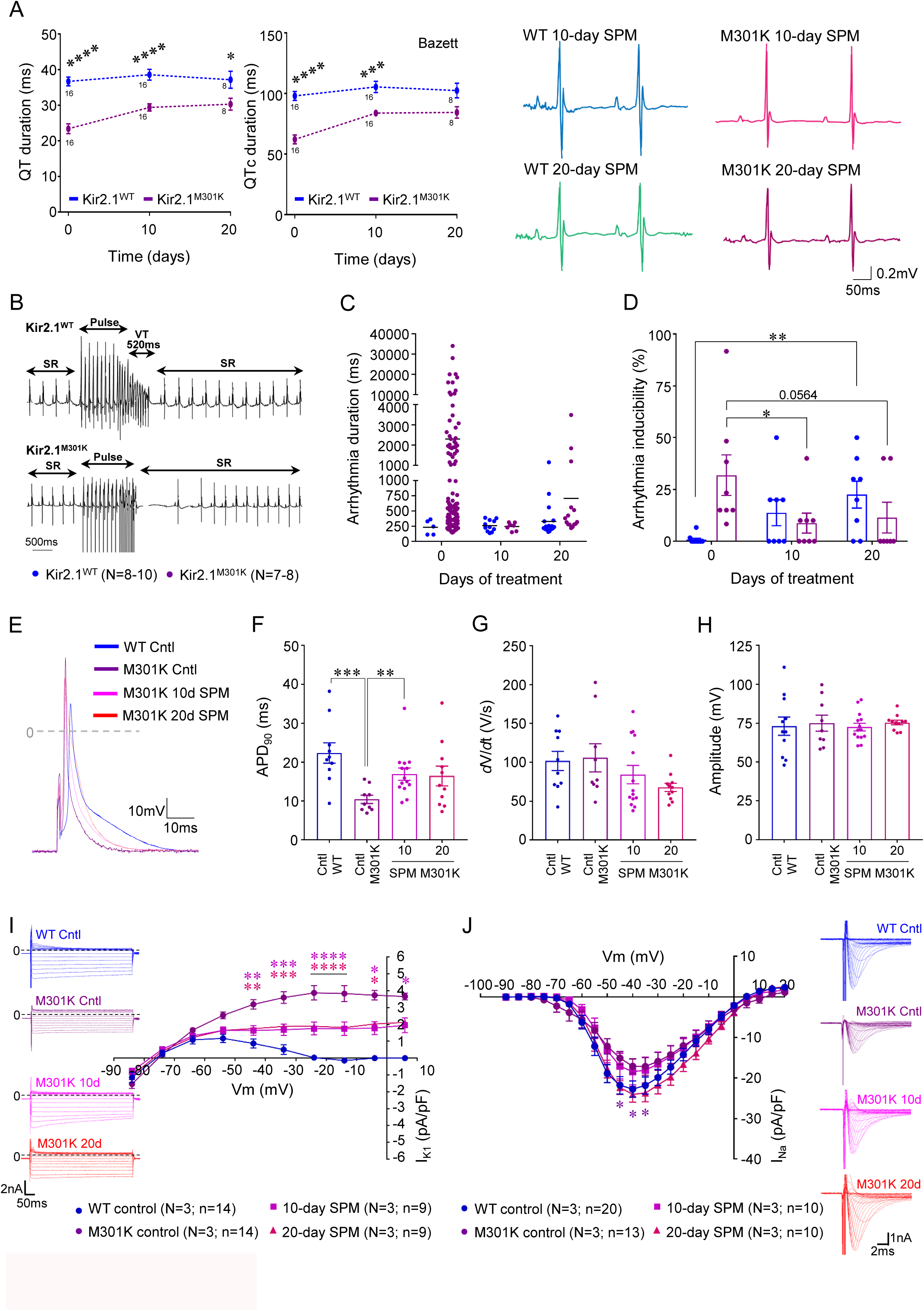
Administration of spermine reduced the arrhythmia inducibility and the I_K1_ hyperfunctionality in Kir2.1^M301K^ mice. **A**, <u>Left</u> and <u>middle</u>, QT and QTc (Bazett’s correction) intervals in Kir2.1^WT^ (blue) and Kir2.1^M301K^ (purple) at baseline and after 10- and 20-day treatment with 10mg/Kg/day spermine (N=16 mice per condition at time 0 and 10-day, and N=8 at 20-day treatment). <u>Right</u>, representative ECG traces after 10- day and 20-day treatment with spermine. **B**, Representative traces of the ventricular arrhythmia recorded after 20 days of treatment. **C**, Duration (ms) of arrhythmias at 0, 10- day and 20-day treatment with spermine in Kir2.1^WT^ (N=10 non-treated and N=8 treated mice) and Kir2.1^M301K^ (N=8 control, N=8 10-day treated, and N=7 20-day treated mice). Note the reduction in number and duration of the arrhythmogenic events in Kir2.1^M301K^ mice. **D**, Arrhythmia inducibility after ventricular electrical stimulation at 0, 10-day and 20-day treatment with spermine. **E**, Representative traces of APs recorded from WT control (dark blue), M301K control (dark purple), M301K 10-day spermine (light purple), and M301K 20-day spermine (dark pink). **F-H**, APD_90_, *d*V/*d*t, and AP amplitude measured from the AP recorded from WT control (N=3; n=10), M301K control (N=3; n=9), M301K 10-day spermine (N=3; n=14), and M301K 20-day spermine (N=3, n=11) ventricular cardiomyocytes, respectively. **I**, Outward I_K1_ recorded from WT control (N=3, n=14), M301K control (N=3, n=14), M301K 10-day spermine (N=3, n=9), and M301K 20-day spermine (N=3, n=9) ventricular cardiomyocytes. **J**, I_Na_ IVs recorded from WT control (N=3, n=20), M301K control (N=3, n=13), M301K 10-day spermine (N=3, n=10), and M301K 20-day spermine (N=3, n=10) myocytes. Spermine is indicated as “SPM” through the panels. Mann-Whitney test and two-way ANOVA (corrected by Šídák’s multiple comparisons) were applied (****p<0.0001, ***p<0.001, **p<0.01, and *p<0.05).

At the cellular level, current-clamp recordings in ventricular cardiomyocytes isolated from spermine-treated Kir2.1^M301K^ mice revealed a progressive APD prolongation compared to untreated Kir2.1^M301K^ (**Figure 6E-H** and **Supplementary Figure 10B**), with no change in the RMP (**Supplementary Figure 10C**). As shown by the superimposed IV relations in **Figure 6I**, chronic i.p. administration of spermine significantly reduced the outward component of I_K1_ in the Kir2.1^M301K^ cardiomyocytes. Over time, spermine also restored the I_Na_ peak current and substantially reduced the Na^+^ window current (**Figure 6J** and **Supplementary Figure 10D**-E).

Spermine treatment did not significantly change the RMP, *d*V/*d*t, APA (**Supplementary Figure 11A**-B), and only slightly prolonged the APD of Kir2.1^WT^ cardiomyocytes (**Supplementary Figure 11C**-D), attributed to a mild reduction in the outward amplitude of I_K1_ (**Supplementary Figure 11E**). Spermine treatment did not functionally alter the peak I_Na_ or its activation/inactivation properties in Kir2.1^WT^ cardiomyocytes (**Supplementary Figure 11F**-H).

Altogether, the results thus far indicate that the systemic administration of 10mg/Kg/day spermine for up to 20 days has beneficial effects in SQTS3 mice expressing Kir2.1^M301K^. It increases I_K1_ rectification prolonging the APD. Consequently, it increases the QT interval and reduces arrhythmia inducibility.

### Chronic oral spermidine administration prolongs the QT interval and reduces arrhythmia inducibility in Kir2.1^M301K^ animals

Oral administration is the most convenient, cost-effective, and commonly used route of drug therapy. Oral administration of spermidine, the second most potent voltage dependent Kir2.1 blocker, is available at the pharmacies as a food supplement.^16, 18, 22–24^ It has been suggested to be cardioprotective and some authors have reported promising results, although in different contexts.^39, 57–61^ Therefore, we decided to add 3mM spermidine to the drinking water of 15- to 17-week-old Kir2.1^WT^ and Kir2.1^M301K^ mice for 20 days (to compare with spermine) and 6 months (for a long-term treatment study). The treatment significantly prolonged the QT/QTc interval over time without significantly altering other ECG parameters in ether Kir2.1^WT^ or Kir2.1^M301K^ mice (**Figure 7A** and **Supplementary Figure 12A**). PR interval was transiently prolonged in Kir2.1^WT^ mice after 20 days of spermidine treatment, but the effect was not sustained over time; and the QRS was shortly reduced in Kir2.1^M301K^ animals (**Supplementary Figure 12A**). Most importantly orally administered spermidine reduced the arrhythmia duration and inducibility in Kir2.1^M301K^ animals (**Figure 7B-D**). Underlying the QT prolongation, the APD was longer, along with an almost complete restoration of I_K1_ function (**Figure 7E-I** and **Supplementary Figure 12B**-C), and a full reversal of I_Na_ defects in Kir2.1^M301K^ ventricular cardiomyocytes following long-term oral spermidine administration (**Figure 7J** and **Supplementary Figure 12D**-E). The electrophysiological effects of spermidine after 20 days of treatment were comparable to those observed after 6 months, meaning that effects plateau after 3 weeks.

**Figure 7.**
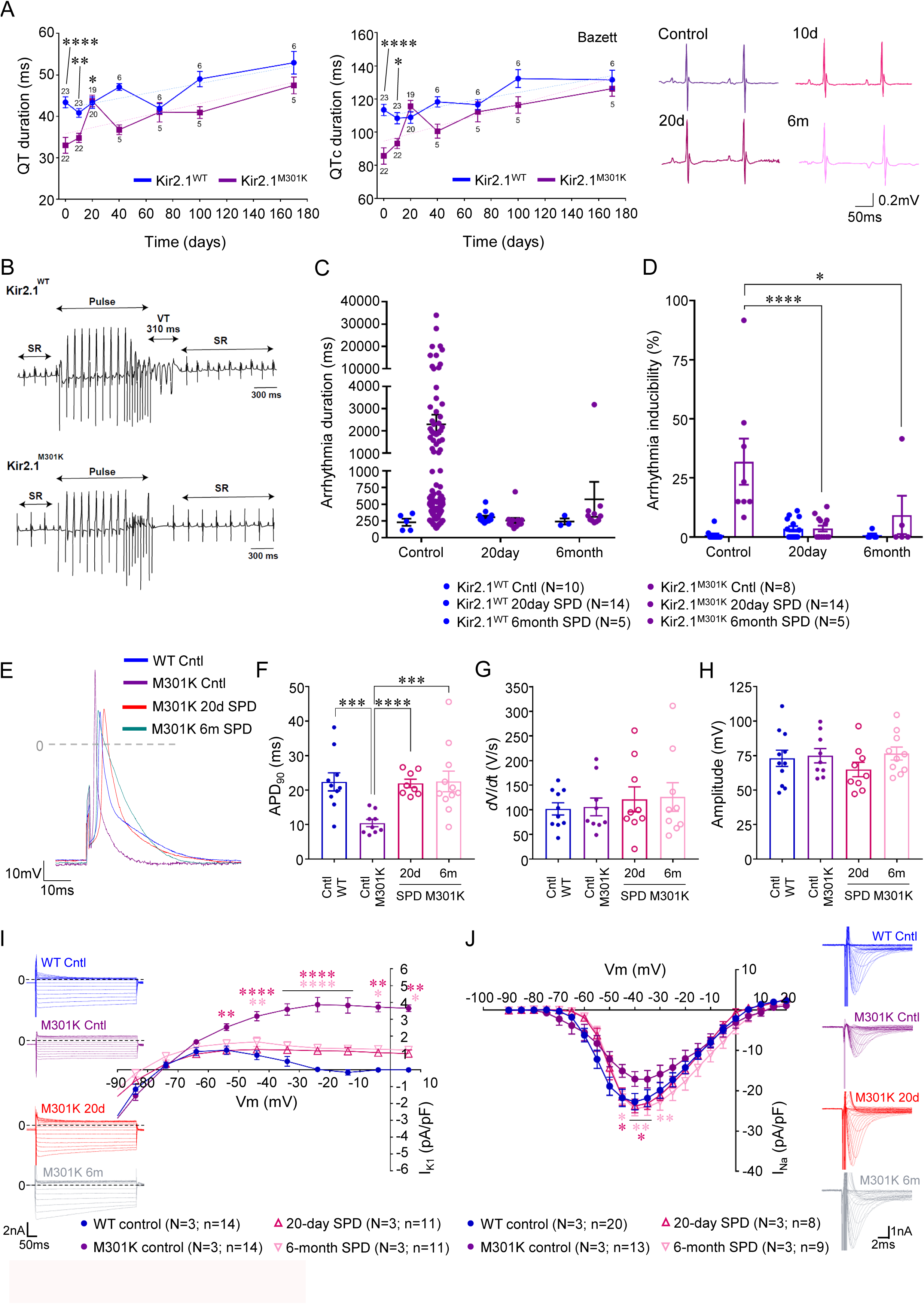
Chronic oral administration of spermidine prolonged the QT interval, reduced the arrhythmia inducibility and ameliorated the I_K1_ and I_Na_ alterations in Kir2.1^M301K^ animals. **A**, <u>Left</u> and <u>middle</u>, QT and QTc (Bazett’s correction) intervals in Kir2.1^WT^ (blue) and Kir2.1^M301K^ (purple) at baseline and over the time after starting the treatment with 3mM spermidine. Sample size is indicated in every time point. <u>Right</u>, representative ECG traces of the same Kir2.1^M301K^ animal untreated (control) and after 10-day, 20-day and 6-month treatment with spermidine via drinking water. **B**, Representative traces of the ventricular arrhythmia recorded after 6 months of treatment. **C**, Duration (ms) of the arrhythmias induced in Kir2.1^WT^ (N=10 control, N=14 20-day treatment, and N=5 6-month treated mice) and Kir2.1^M301K^ (N=8 control, N=14 20-day treated, and N=5 6-month treated mice). Note the reduction in number and duration of the arrhythmogenic episodes in Kir2.1^M301K^ mice. **D**, Arrhythmia inducibility of Kir2.1^WT^ and Kir2.1^M301K^ mice after ventricular pacing at different days of treatment. **E**, Representative traces of APs recorded from WT control (dark blue), M301K control (dark purple), M301K 20-day spermidine (dark pink), and M301K 6-month spermidine (light pink). **F-H**, AP parameters: APD_90_ (**F**), *d*V/*d*t (**G**), and AP amplitude (**H**) measured from WT control (N=3; n=10), M301K control (N=3; n=9), M301K 20-day (N=3; n=8-9), and M301K 6-month spermidine (N=3, n=9-11) ventricular cardiomyocytes. **I**, Outward I_K1_ recorded from WT control (N=3, n=14), M301K control (N=3, n=14), M301K 20-day spermidine (N=3, n=11), and M301K 6-month spermidine (N=3, n=11) ventricular cardiomyocytes. **J**, I_Na_ IVs recorded from WT control (N=3, n=20), M301K control (N=3, n=13), M301K 20-day spermidine (N=3, n=8), and M301K 6-month spermidine (N=3, n=9) myocytes. Spermidine is indicated as “SPD” through the panels. Mann-Whitney test and two-way ANOVA corrected by Šídák’s multiple comparisons test were applied (****p<0.0001, ***p<0.001, **p<0.01, and *p<0.05).

Additionally, spermidine did not modify the RMP, *d*V/*d*t or APA (**Supplementary Figure 13A**-B), but it did prolong significantly the APD_90_ in Kir2.1^WT^ cardiomyocytes (**Supplementary Figure 13C**-D). Underlying this prolongation, was a reduction in the outward amplitude of I_K1_ both 20-day and 6-month after starting spermidine administration (**Supplementary Figure 13E**). The peak I_Na_ of Kir2.1^WT^ cardiomyocytes was unchanged by spermidine, although it slightly shifted toward depolarized voltages (**Supplementary Figure 13F**-H).

### Oral spermidine exhibits a favorable safety profile while maintaining outward I_K1_ reduction in long-term treated mice

Young (16-week-old) and old (55-week-old) Kir2.1^WT^ and Kir2.1^M301K^ animals under long-term (6-month) oral treatment with 3mM spermidine underwent functional, structural, and serological analyses to determine drug toxicity. Similar to young adult mice, aged Kir2.1^M301K^ animals exhibited prolonged QT/QTc intervals over time following the initiation of long-term spermidine treatment (**Supplementary Figure 14**). After 3 weeks, these values reached levels comparable to those of Kir2.1^WT^ mice, without significant disturbances in any other ECG parameter. In contrast, untreated Kir2.1^WT^ and Kir2.1^M301K^ old mice—littermates of those receiving 3mM spermidine for 6 months—displayed shortened QT/QTc intervals, consistent with the pattern seen in young adults (**Supplementary Figure 14A**-B). Like with spermine, both groups showed a body weight reduction at 6 months spermidine treatment (**Supplementary Figure 15A**-B), but the body weight/heart weight ratio did not change compared to untreated untransfected control animals (**Supplementary Figure 15C**). Echocardiography showed no differences in ejection fraction or fractional shortening between untreated and long-term treated mice, either young or old adult (**Figure 8A-B**). Histological analysis following long-term treatment in young animals revealed no structural abnormalities or evidence of fibro-adipose replacement in the liver, kidney, lung or spleen (**Supplementary Figure 15D**). Additionally, the hearts from treated mice exhibited normal structure, comparable to those from untreated untransfected control animals (**Figure 8C**). Serological analyses and biochemical parameters measured in the same young adult mice were within normal ranges (**Supplementary Figure 16**).^62, 63^ Together, these results demonstrate that chronic oral spermidine administration is safe and effective, means to restore the QT interval and prevents arrhythmia inducibility in our SQTS3 mouse model.

**Figure 8.**
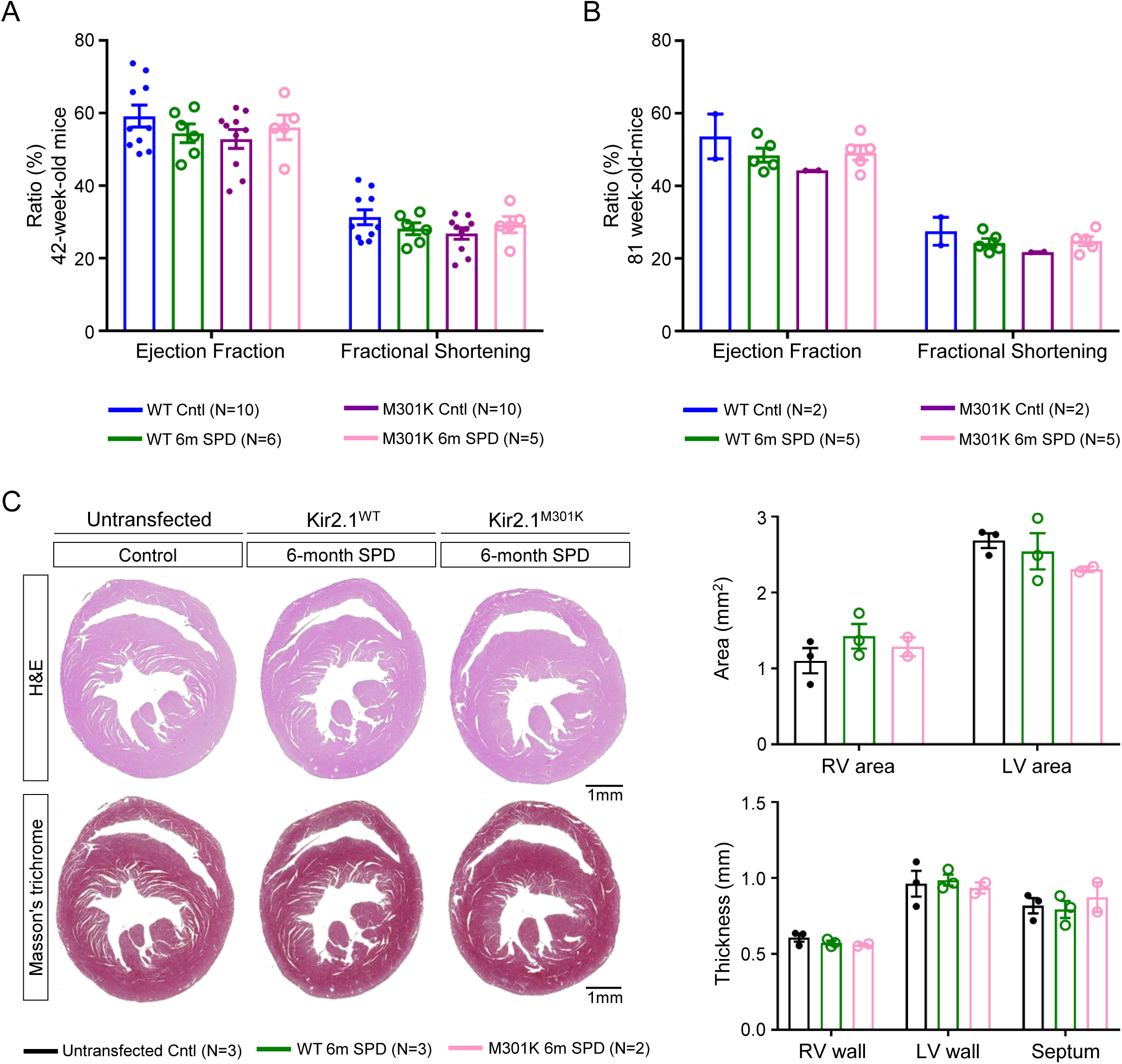
Oral long-term treatment with spermidine exhibits safe and nontoxic profile in either Kir2.1^WT^ or Kir2.1^M301K^ mice. **A-B**, Echocardiography analysis showing ejection fraction and fractional shortening of untreated and 6-months spermidine treated Kir2.1^WT^ and Kir2.1^M301K^ young adult (42-week-old) (**A**) and old adult (81-week-old) (**B**) mice. Kir2.1^WT^ untreated are shown in dark blue (N=2-10), Kir2.1^M301K^ untreated in dark purple (N=2-10), Kir2.1^WT^ treated in dark green (N=5-6), and Kir2.1^M301K^ treated in light pink (N=5). **C**, Haematoxylin and Eosin (H&E), and Masson’s trichrome staining of untreated untransfected and 6-months treated Kir2.1^WT^ and Kir2.1^M301K^ hearts. Top, The right ventricle (RV) and left ventricle (LV) areas are presented. Bottom, Graphs show quantification of the RV, LV, and septum wall thickness (N=2-3 animals per condition). Spermidine is indicated as “SPD” through the panels. Two-way ANOVA tests were applied for comparisons.

## DISCUSSION

Currently, the absence of mechanistically targeted therapies for SQTS3 is in part due to the unknown mechanisms underlying some of the identified mutations. We aimed to decipher the arrhythmogenic and molecular mechanisms that make the human Kir2.1^M301K^ mutation causing the most abbreviated QT interval described in the literature^11^ with the final goal of enhancing the effective therapeutic approaches in SQTS3. Heterologous expression systems are too limited for this; hence we generated the first *in-vivo* SQTS3 mouse model carrying the same variant, using Kir2.1^WT^ as a control. Introduction of exogenous human Kir2.1^M301K^ in mice expressing the endogenous murine Kir2.1 reproduces a heterozygous condition where the dominant negative effect of the mutation mimics the patient’s genetic environment in the heart. Importantly, we confirmed that transduction using the human WT or mutant Kir2.1 sequences did not cause neither Kir2.1 overexpression nor structural alterations in the cardiac tissue (**Supplementary Figure 1**).

Our generated AAV9-mediated mouse model presented an extremely shortened QT interval and long-lasting ventricular arrhythmias induced by intracardiac stimulation, mirroring the patient’s electrophysiological phenotype. Kir2.1^M301K^ cardiomyocytes had extremely abbreviated APD, linked to impaired inward rectification and increased I_K1_ at voltages above -80 mV. Amino acid M301 is close to E299 in the Kir2.1 channel structure, and polyamines bind to this region to block the channel at depolarized voltages.^64, 65^ The replacement of methionine (M) with lysine (K)―a polar, positively charged amino acid―at position 301 electrostatically impairs polyamine binding. This results, on the one hand, in a gain of Kir2.1 function due to reduced inward-going rectification when expressed in heterozygosis. Heterotetrameric Kir2.1^WT^-Kir2.1^M301K^ channels remain open at voltages where they would normally be closed, leading to a gain-of-function. On the other hand, the addition of positively charged lysine residues in the cytoplasmic pore of Kir2.1 interferes the potassium conductance. Consequently, when Kir2.1^M301K^ is expressed in homozygosis, the resulting mutant homotetrameric conformation of Kir2.1―with 4 lysines in the cytoplasmic pore―electrostatically repels the potassium flow through the channel and leads to a loss-of-function. Therefore, the electrophysiological consequences of Kir2.1^M301K^ differ depending on whether it is expressed in homo- or heterozygous conditions. While in the homozygous condition the mutation reduces I_K1_, it dramatically increases the current when co-expressed with Kir2.1^WT^.^11^

While analyzing the ECG from Kir2.1^M301K^ mice, we realized that the QRS complex was significantly prolonged compared to Kir2.1^WT^ animals. Unfortunately, we did not have access to the ECGs of the young girl carrying the Kir2.1^M301K^ mutation before her sudden demise and have not been able to determine whether her QRS complex was prolonged. To study the possible reason for the QRS prolongation caused by Kir2.1^M301K^ in the mouse, we isolated ventricular cardiomyocytes and demonstrated that this genetic variant disturbs Na_V_1.5 function by reducing the I_Na_ density and modifying the I_Na_ activation curve at negative voltages. The latter effect increased the window current and predisposed the heart to premature ventricular contractions and tachycardia, as has been described for the I151V mutation in Na_V_1.5, among others.^66^ Therefore, quinidine may be contraindicated in these patients.

Kir2.1 and Na_V_1.5 work synergistically, and loss-of-function mutations in Kir2.1 reduce the Na_V_1.5 current density and *vice versa*.^45–47, 50, 54, 55^ Recently, we demonstrated that the SQTS3-related mutation Kir2.1^E299V^ modified I_Na_ in a cardiac chamber-specific manner.^51^ Moreover, *in-silico* simulations predicted that both channels have a physical interaction interface with a favorable binding energy: the voltage domains DII-S1 of Na_V_1.5 interact with the transmembrane helix M1 of Kir2.1.^67^ Kir2.1 has been shown to interact with as many as 218 high-confidence interactors encompassing various molecular mechanisms of Kir2.1 function, ranging from intracellular trafficking to lysosomal degradation.^48^ In a different study, membrane-enriched cardiac lysates of male mice, identified 61 protein interactors for *KCNJ2* and 59 for *SCN5A*.^68^ Kir2.1 was not listed as an interactor of *SCN5A*, and Na_V_1.5 was not among the detected interactors for *KCNJ2*. However, both channels share some common proteins, such as apolipoproteins, polypeptides and E3 ubiquitin-protein ligases,^68^ hence a third partner could mediate the functional relationship between them. Proteins like SAP97 and α1- syntrophin act as scaffolds that keep Kir2.1 and Na_V_1.5 together at the membrane through their PDZ binding domains. *In-vitro* experiments have demonstrated that inhibition or absence of these interacting proteins causes alterations in both I_K1_ and I_Na_.^45, 69–74^ However, we have not seen any changes in the distribution of these specific interactors in ventricular cardiomyocytes expressing the Kir2.1^M301K^ mutation. Unlike SAP97 and α1-syntrophin, we did see a reduction in the total and sarcolemmal protein levels of Na_V_1.5 in Kir2.1^M301K^ ventricular cardiomyocytes, so other unknown common interactors might mediate, promote, or even enhance these alterations. Hattori T *et al*. demonstrated that the homozygous expression of Kir2.1^M301K^ in HEK cells resulted in an absence of I_K1_,^11^ so the I_Na_ reduction seen in heterozygous cardiomyocytes could be the result of a destabilization of Na_V_1.5 when it is in contact with Kir2.1^M301K^ homotetrameres.

While the molecular basis of how Kir2.1^M301K^ reduced I_Na_ is beyond the scope of this study, one possible explanation is that the mutation may affect the interaction of both channels with 14-3-3 proteins and indirectly alter Na_V_1.5 phosphorylation.^55, 75–77^ Alternatively, Na_V_-β subunits could mediate the Kir2.1^M301K^ effects, as they form multiprotein complexes with ion channels and are essential for Na_V_1.5 function in cardiomyocytes.^78–84^ Yet another explanation is that Kir2.1^M301K^ may alter Na_V_1.5 degradation, as both channels share some E3 ubiquitin-protein ligases.^68^ However, further research is needed to decipher how the mutation Kir2.1^M301K^ reduces the expression and current density of Na_V_1.5 channels.

On the other hand, in this work we also aimed to increase the mechanistically targeted treatment options for SQTS3 patients. Since the pathogenic mechanism of the Kir2.1^M301K^ mutation involves a loss of inward rectification, and polyamines can bind to the Kir2.1 cytoplasmic pore to block the channel in a voltage-dependent manner,^23, 85^ we sought to evaluate the effects of their chronic administration. It has been previously shown that polyamine administration is cardioprotective and reportedly has anti-aging effects.^39, 57–61^ They are also effective in neurodegenerative diseases by inducing autophagy, oxidative stress, and amyloid-β accumulation.^86^ In hypertension, heart failure, and myocardial infarction, polyamines have been shown to promote cardiac autophagy and longevity, and to reduce inflammation and oxidative damage.^87, 88^ In metabolic disorders, such as type 2 diabetes, obesity, and fatty liver disease, they improve insulin sensitivity, lipid metabolism, and reduce hepatic steatosis.^89^ In different cancers, they enhance anti-tumour immune responses;^90^ and in inflammatory and autoimmune diseases like rheumatoid arthritis and inflammatory bowel disease, polyamines reduce pro-inflammatory cytokines and promote intestinal barrier integrity.^91^

In this sense, we recently demonstrated that increasing the cytoplasmic concentration of spermine interfered with the I_K1_ gain-of-function of some mutant Kir2.1 channels that lacked inward-going rectification.^51^ Therefore, we hypothesized that repurposing polyamines by administering them to control hyperfunctional Kir2.1^M301K^ channels would have therapeutic benefit. Spermine is the polyamine with the most powerful capacity for blocking Kir2.1 channels,^16, 18, 24^ but spermidine, the second most potent blocker of Kir2.1, is already available as a food supplement, having demonstrated beneficial effects at different levels (see above). We administered either 10mg/Kg/day spermine i.p. or 3mM spermidine via drinking water and, as a checkpoint to be sure that both treatments were having effect, we noticed that all animals lost about 7.5% of their weight after 10 days of treatment, since it has been described that polyamines improve glucose metabolism in mice.^39^ Most importantly, we demonstrated that chronic administrations of spermine or spermidine prolonged the APD and the QT interval, and reduced the arrhythmia incidence in Kir2.1^M301K^ mice. Using untransfected, untreated mice of the same age as controls in some experiments, we also shown that neither AAVs, Kir2.1^M301K^ expression, nor long-term SPD treatment altered cardiac function, structure, or serological profiles. Moreover, polyamines reversed QRS prolongation by restoring the I_Na_ defects caused by the Kir2.1^M301K^ mutation. Further research is needed to understand the mechanisms underlying these effects, but it seems that the Kir2.1^M301K^-derived I_Na_ alterations depend on the I_K1_ gain-of-function at depolarized potentials. When this gain-of-function is alleviated, the I_Na_ disturbances provoked by Kir2.1^M301K^ are similarly reduced. Interestingly, polyamines were shown to prolong the QT interval in mutant mice regardless of whether treatment began at early or late stages of aging.

The absence of arrhythmic events in most polyamine-treated mutant mice is highly promising; polyamine treatment reduces the arrhythmogenic potential of a highly pathogenic mutation in mice, which opens a new pathway for investigation at the preclinical and clinical levels. Therefore, based on our findings, we conclude that chronic treatment with polyamines could be considered a potential therapeutic strategy for patients with SQTS3. Targeting the underlying electrophysiological abnormalities of Kir2.1^M301K^, using polyamines or novel derivatives, may help prevent life-threatening arrhythmias and reduce the risk of SCD in these patients.

## LIMITATIONS

Data derived from our AAV9-mediated cardiac-specific mouse model of SQTS3 should not be directly extrapolated to the human disease and should be cautiously interpreted. First, heart rate and repolarization parameters differ substantially between mice and humans, as different groups of ionic currents govern the ionic mechanisms. There are also remarkable structural differences between murine and human hearts, particularly in heart size. Therefore, additional studies using appropriate preclinical models would be needed to ensure rigorous translation of our results to the patient. Second, although cardiac-specific AAV9 vectors delivered intravenously show 95% of transduction rate and over 50% of the cardiomyocytes take between 2 and 4 copies of the transgene, this system always generates a mosaic cellular distribution of mutant and wildtype expression throughout the heart. However, this mosaicism is also present in AAV9-Kir2.1^WT^, the condition used as a control. Yet, despite the above limitations, we stand by our results, which provide novel insights into mechanisms underlying this arrhythmogenic and potentially lethal channelopathy and into the innovative role that polyamines could take in the treatment of SQTS3 patients.

## DATA AVAILABILITY STATEMENT

The data underlying this article are available in the article and in its online supplementary material. Additional data will be shared on request to the corresponding author.

## AUTHOR CONTRIBUTION

A.I.M.M and J.J. co-designed the experiments; A.I.M.M., A.M. and F.M.C. performed all the experiments contained in that paper; A.I.M.M characterized *ex vivo* the AAV9 infection level of WT and SQTS3 mouse hearts, studied their electrical phenotype *in vivo*, isolated cardiomyocytes, performed most of the electrophysiological experiments, and carried out the molecular and cellular biology techniques, along with their analysis; F.M.C. infected the C57BL/6 mice using the appropriate AAV9 and contributed with the mouse models *in-vivo* characterization after treating the mice with polyamines; A.M. conducted some of the cellular electrophysiology experiments, both before and after starting the treatment with spermine, and provided some funding; E.C.B. analyzed the echocardiography; A.I.M.M. and J.J. co-wrote the manuscript and conceived the study; J.J. provided supervision, funding and revisions; all authors discussed the results and approved the manuscript.

## ACKNOWLEDGEMENTS

We thank the Cardiac Arrhythmia Laboratory members, with a special mention to Dr. Patricia Sánchez-Pérez and Dr. Paula G. Socuéllamos, in addition to Dr. Carlos Morillo, for providing discussions and revisions. We thank the CNIC Viral Vectors Unit for producing the AAV9 used in this article. The confocal experiments were carried out in the CNIC Microscopy and Dynamic Imaging Unit – ICTS-ReDib with funding from MCIN/AEI /10.13039/501100011033 and FEDER “Una manera de hacer Europa” (#ICTS-2018-04-CNIC-16).

## SUPPLEMENTAL INFORMATION

Extended Materials and Methods Supplementary Tables 1-8 Supplementary Figures 1-16 References 1-13

