## Supplementary Material for "Repurposing polyamines to prevent arrhythmias in Short QT Syndrome type 3, a potentially lethal disease"

### Repurposing polyamines for the treatment of Short QT Syndrome type 3, a potentially lethal disease

[1] Centro Nacional de Investigaciones Cardiovasculares (CNIC), 28029, Madrid, Spain.
[2] Cardiology Department, Virgen de las Nieves University Hospital, 18014, Granada, Spain.
[3] Ibs. Granada Instituto de Investigación Biosanitaria, 18012, Granada, Spain.
[4] CIBER de Enfermedades Cardiovasculares (CIBERCV), Madrid, Spain.
[5] Departments of Internal Medicine and Molecular and Integrative Physiology, University of Michigan, 48109, Ann Arbor, MI, USA.

* These authors contributed equally.

[#]Corresponding author:

José Jalife, MD, PhD
Distinguished Senior Investigator Centro Nacional de Investigaciones Cardiovasculares (CNIC) Carlos III
Melchor Fernández Almagro, 3, 28029, Madrid, Spain
Emeritus Professor of Internal Medicine, University of Michigan, Ann Arbor, MI, USA


**Short title:** Repurposing polyamines to treat SQTS3 patients.

### EXTENDED MATERIALS AND METHODS

**Study approval.** All experimental and other scientific procedures using animals conformed to EU Directive 2010/63EU and Recommendation 2007/526/EC, enforced in Spanish law under *Real Decreto 53/2013*. The local ethics committees and the Animal Protection Area of the Comunidad Autónoma de Madrid (PROEX 226.5/23) approved all animal protocols.

**Mice.** 4-week-old C57BL/6J male mice were obtained from Charles River Laboratories. Mice were reared and housed following institutional guidelines and regulations. The mice had access to food and water *ad libitum*. Fifteen- to 35-week-old mice were used for experiments when we needed untreated or short-term treated animals. For long-term spermidine treatment, we used 16-week-old and 55-week-old Kir2.1$^{WT}$ and Kir2.1$^{M301K}$ mice, and studied their electrophysiological phenotype for up to 6 months after starting treatment (i.e., when they were 42 and 81 weeks old, respectively). Untransfected untreated animals were used as controls when indicated. Experimental procedures were conducted in anesthetized animals, which is known to pause the circadian clock.[1] Nevertheless, experiments in which the circadian rhythm could be relevant (ECGs, echocardiography, cardiomyocyte isolation, and intracardiac stimulation) were done during daylight hours, specifically, from 9 am to 2 pm.

**AAV production and purification.** Briefly, this was achieved by first generating enhanced tdTomato-reporter AAV vectors driven from the cardiomyocyte-specific cardiac Troponin T proximal promoter (cTnT), strengthened by a *cis*-regulatory motif number 4 enhancer (CRM4),[2] and encoding wildtype Kir2.1 (Kir2.1$^{WT}$), or the SQTS3 Kir2.1 mutant (Kir2.1$^{M301K}$). These AAV plasmids were packaged into AAV serotype 9 (AVV9) capsids with the use of pAdDF6 helper plasmids (providing the three *adenoviral helper* genes) and pAAV2/9 (providing *rep* and *cap* viral genes), obtained from PennVector by the CNIC Viral Vectors Unit.[3] AAV vectors were produced by the triple transfection method, using HEK-293A cells as described previously.[4, 5] AAV vector titer (viral genomes, vg, per ml) was carried out by quantitative real-time PCR (qRT-PCR) as described.[6] Known copy numbers ($10^5$–$10^8$) of the respective plasmid (pAAV-empty vector, pAAV-Kir2.1$^{WT}$, and pAAV-Kir2.1$^{M301K}$) were used to construct standard curves.

**AAV injection.** Mice were anesthetized with 60mg/kg Ketamine and 20mg/kg Xylazine administered intraperitoneally (i.p.). Once asleep, animals were placed on a heated pad at 37±0.5ºC to prevent hypothermia. Thereafter, we administered $3.5 \times 10^{10}$ vg to 4- to 5-week-old mice intravenously (i.v.) via the femoral vein in a final volume of 50μL, taking care to prevent the introduction of air bubbles. Animals were then maintained on the heating pad until recovery.

**AAV-mediated gene distribution.** We assessed the corporal distribution of AAV-mediated protein expression eight weeks after AAV9 administration.[3] The *ex-vivo* fluorescent signal was detected in hearts from transfected mice confirming the cardiac expression of protein tdTomato used as the reporter gene in the viral construct. The infection rate in mouse hearts (measured as vg/cell) was quantified blindly by immunohistochemistry using an anti-tdTomato antibody and a home-made Fiji plugin, as previously described.[3]

**Echocardiography.** Mice were lightly anesthetized with 0.5-2% isoflurane in oxygen, and placed on a 37ºC heating platform in the supine position. A base-to-apex electrocardiogram (ECG) continuously monitored the animal, adjusting isoflurane delivery to maintain the heart rate at 450±50 beats per minute (bpm). Warmed ultrasound gel was used to maintain normothermia. Transthoracic echocardiography was performed blindly using a high-frequency ultrasound system (Vevo 2100, Visualsonics Inc., Canada) with a 40-MHz linear probe. Two-dimensional (2D) guided M-mode echocardiograms were obtained at a frame rate above 230 frames/s, and pulse wave Doppler was acquired at a frequency of 40kHz. Images were transferred to a computer and analyzed blindly offline using the Vevo 2100 Workstation software. The M-mode sample gate was placed at the level of the papillary muscles. Parasternal standard long and short axis views were acquired to assess left ventricular (LV) systolic function. LV ejection fraction (EF) and chamber dimensions were calculated from these views in systole and diastole.

**Surface ECG recordings.** Mice were anesthetized adjusting isoflurane (1.5-2% volume in oxygen) delivery to maintain the heart rate at 350±50bpm and the efficacy of the anesthesia was controlled by monitoring the breathing rate and the absence of response to painful stimuli. A subcutaneous 23-gauge needle electrode connected to an MP36R amplifier (BIOPAC Systems) was attached to each limb, and six-lead surface ECGs were recorded.

We used AcqKnowledge 4.1 software and LabChart 8 Reader to analyze, blindly and, sometimes in pairs, the ECG complexes, segments and interval durations. For each animal, we measured the duration of these parameters in 5 to 10 beats per time of interest, and the mean ± standard error of the mean (SEM) was presented. We applied Bazett's and Framingham's formulas to correct the QT interval considering the heart rate.

$$QTc\ (Bazett) = \frac{QT}{\sqrt{RR}} \qquad QTc\ (Framingham) = QT + 0.154\ (1 - RR)$$

**Polyamine administration.** First, 15- to 17-week-old male mice were used to test the effects of polyamine administration. Following previously published indications, 10mg/Kg spermine (Sigma-Aldrich, 71-44-3), was administered i.p. daily for 10 or 20 days, changing the administration area to avoid inflammation.[7] In addition, 3mM spermidine (Sigma-Aldrich, S2626) was administered via drinking water for 10 or 20 days, or a maximum of 6 months, changing the water and preparing fresh spermidine every 3 days.[8] Fifty-two-week-old mice also underwent long-term treatment with spermidine.

***In-vivo* intracardial electrophysiological studies.** After anesthesia (60mg/kg Ketamine and 20mg/kg Xylazine i.p.), an octopolar catheter (Science) was inserted through the jugular vein and advanced into the right atria and right ventricle.[9, 10] The animal was placed on a 37ºC heating platform in the supine position.

To measure the duration of the refractory period, we used an S1-S2 protocol consisting of a train of 10 stimuli (S1) at a cycle length of 100ms (10Hz) followed by an extra-stimulus (S2) reducing the S1-S2 interval in 2-ms steps starting at 100ms. Atrial and ventricular arrhythmia inducibility was assessed by applying 10 stimuli at 10Hz (S1), followed by 10 stimuli at 25Hz (S2), all of them 2ms (duration) and 7V (amplitude)-stimuli. Atrial fibrillation was defined as the occurrence of rapid and fragmented atrial electrograms (lack of P waves) with irregularly irregular atrio-ventricular (AV)-nodal conduction and ventricular rhythm. Ventricular tachycardia/ventricular fibrillation (VT/VF) was defined as an exceedingly fast monomorphic/polymorphic ventricular rhythm independent of atrial excitation. Frequency of the recorded VT episodes lasting more than 500ms were measured as the peaks per second.

**Cardiomyocyte isolation.** To isolate ventricular cardiomyocytes we used the procedure of Macías *et al.*[11] Briefly, after cervical dislocation, the mouse heart was isolated, mounted on a modified Langendorff-perfusion apparatus and the aorta was retrogradely perfused with $Ca^{2+}$-free perfusion buffer (PB) containing (in mmol/L): 113 NaCl, 4.7 KCl, 0.6 $KH_2PO_4$, 0.6 $Na_2HPO_4$, 1.2 $MgSO_4 \cdot 7H_2O$, 12 $NaHCO_3$, 10 $KHCO_3$, 0.032 Phenol Red, 9.8 HEPES, 30 taurine, 5.5 glucose, 10 2,3-butanedione-monoxime (pH 7.4 with NaOH). Perfusion rate was 1mL/min for 7min at 37°C. Enzymatic digestion was then started for 20min with digestion buffer (DB) consisting of PB supplemented with Liberase™ (0.2mg/mL), Trypsin 2.5% (5.5mmol/L), and $CaCl_2$ (12.5µM) at 37°C. Thereafter, ventricular cardiomyocytes were gently disaggregated and isolated in 3mL DB. The resulting cell suspension was filtered through a 200µm sterile mesh (SEFAR-Nitex) and transferred for enzymatic inactivation to a tube containing 12ml stopping-buffer-1 (SB-1), consisting of PB supplemented with fetal bovine serum (FBS, 10% v/v) and $CaCl_2$ (12.5µmol/L). After gravity sedimentation for 20min, the supernatant was removed and cardiomyocytes were resuspended in stopping-buffer-2 (SB-2) containing the same concentration of $CaCl_2$, but lower FBS (5% v/v) for another 20min. Cardiomyocyte $Ca^{2+}$-reintroduction was performed in SB-2 with two progressively increased $CaCl_2$ concentrations (0.112 and 1mmol/L). Cells were resuspended and allowed to decant for 15min in each step, contributing to the purification of the cardiomyocyte suspension.

**Patch-clamping of isolated cardiomyocytes**. For the patch-clamp experiments in the whole-cell configuration, we used the external (bath) and internal (pipette) solutions that are listed in **Supplementary Table 1**.[11-13]

Cardiomyocytes were placed in a superfusion chamber (RC-26, Warner Instruments) mounted on the stage of an inverted microscope (DMi8, Leica). Cells were allowed to settle on the bottom of the superfusion chamber before being perfused with the corresponding solution. After sealing the membrane, whole-cell voltage-clamp or current-clamp recordings were made using an Axopatch 200B Amplifier (Axon Instruments) and a MultiClamp 700B Microelectrode Amplifier (Molecular Devices). Pipettes made from borosilicate glass (GD-1, Narishige, OD: 1mm; ID: 0.6mm) had resistances of 2-5 MΩ after using the internal solutions. Series resistance compensation of 80-90% was achieved. All voltage-clamp currents were low-pass filtered at 2kHz with an analog filter and digitized at 4-10Hz. Data were recorded using pClamp 10.6 software with the Clampex 10.6 program and analyzed with the Clampfit 10.6 program (Axon Instruments, Foster City, USA). Current amplitudes were normalized to the cell capacitance to account for differences in cell size and expressed as current densities (pA/pF).

<u>Whole-cell current-clamp recordings</u>: We patch-clamped the cardiomyocytes in the current clamp configuration using a conventional patch-clamp amplifier to record changes in the voltage through the membrane ($V_m$). Threshold current was determined using 1ms pulses at increasing amplitudes (0.2nA/pulse) and frequency of 1Hz. Thereafter, action potentials (APs) were evoked by the injection of 1ms pulses of constant amplitude. We recorded AP stimulated at 1, 2, 4, 5 and 10 Hz. APD was measured at 20, 50, 70 and 90 % of repolarization, and, as part of the AP characterization, we also analyzed the resting membrane potential (RMP), the AP amplitude (APA), and the AP upstroke velocity measured as $dV/dt$. In these experimental conditions, the liquid junction potential correction applied was 13.4mV.

**Supplementary Table 1. External and internal solutions used in patch-clamp experiments.**

| | $I_{K1}$ / AP | | $I_{Na}$ | |
|---|---|---|---|---|
| | Bath solution (mM) | Pipette solution (mM) | Bath solution (mM) | Pipette solution (mM) |
| **Reagent** | pH 7.4 (NaOH) | pH 7.2 (KOH) | pH 7.35 (CsOH) | pH 7.2 (CsOH) |
| $CaCl_2$ | 0.9 | 1 | 1 | - |
| CsCl | - | - | 132.5 | - |
| CsF | - | - | - | 135 |
| EGTA | - | 10 / - | - | 10 |
| Glucose | 10 | - | 11 | - |
| HEPES | 5 | 5 | 20 | 5 |
| K-Aspartate | - | 110 | - | - |
| $Na_2ATP$ | - | 4 | - | - |
| KCl | 5.4 | 20 | - | - |
| MgATP | - | - | - | 5 |
| $MgCl_2$ | - | 1 | 1 | - |
| $CdCl_2$ | - | - | 0.1 | - |
| NaCl | 136 | 8 | 5 | 5 |
| $KH_2PO_4$ | 1.2 | - | - | - |

Whole-cell voltage-clamp recordings. For these experiments, we changed the patch-clamp mode from current-clamp to voltage-clamp, where we used a feedback amplifier to apply voltage steps and measure the current needed to maintain the membrane potential at any given level. This enabled us to construct IV relationships for $I_{K1}$ and $I_{Na}$, as follows:

- *Potassium currents*. IV relationship for $I_{K1}$ densities were constructed by applying 500-ms voltage-clamp steps in 10mV increments from -140 to 0 mV from a holding potential of -80mV at a frequency of 0.1Hz at room temperature (RT). $I_{K1}$ was calculated by subtracting currents recorded in the absence and presence of 500μM $BaCl_2$. In these experiments, the liquid junction potential was 13.4mV; it was applied when calculating and presenting the IVs.
- *Sodium currents*. To record $I_{Na}$, cells were held at -120mV and stepped for 100ms from -90 up to +40 mV in 5mV increments at 0.33Hz. $I_{Na}$ was measured at the peak of the current trace. To analyze $I_{Na}$ inactivation, cells were held at -120mV and stepped for 100ms from -140mV up to -20mV in 10mV increments at 0.2Hz. Leak currents were subtracted using the P/4 protocol. Steady-state activation and inactivation curves were fitted by using a Boltzmann equation, in which $V_{50}$ is the half-maximum (in)activation potential and $k$ is the Slope factor.

**Immunohistochemical procedure.** We analyzed the amount of AAV9 infection in mouse hearts (see the “AAV-Mediated Gene Distribution” section). After perfusing the heart with PBS, it was fixed in 4% paraformaldehyde (PFA) in PBS for 48 hours and conserved in 70% ethanol. Samples were mounted in OCT embedding compound and frozen at -20 to -80 °C. We cut 5-7μm thick tissue sections using a cryostat and thaw-mounted the sections onto gelatin-coated histological slides. After permeating (0.2% Triton X-100 in PBS) and blocking (10% normal goat serum (NGS) in PBS), we used the goat polyclonal anti-tdTomato antibody (Sicgen, AB8181-200) overnight (O/N) and the anti-goat HRP or Alexa-488 secondary antibodies for 1 hour at RT. We also incubated cardiac samples, in addition to liver, kidney, lung, and spleen tissues slices, with Hematoxylin-Eosin to study possible structural changes in infected compared with untransfected hearts, and in non-treated *vs* treated with 3mM spermidine for 6 months. Moreover, we used Masson's trichrome staining to measure the percentage of fibrosis in the same samples and conditions.

**Conventional confocal microscopy experiments.** Isolated cardiomyocytes were washed using PBS and fixed for 10min with 4% PFA in PBS. Cardiomyocytes were permeabilized using Triton X-100 0.2% in PBS for 10min and then blocked in suspension with 10% NGS in a solution containing Triton X-100 0.1% in PBS for 1 hour at RT. We incubated O/N the cells with primary antibodies at 4ºC and secondary antibodies (1:500 in all cases) for 1 hour at RT (the antibodies and their dilutions are specified in **Supplementary Tables 2** and **3**). We washed the samples three times after each antibody incubation with PBS. Samples were mounted in Fluoroshield™-DAPI imaging medium (F6057, Merck). Images of individual cardiomyocytes were acquired with a Leica SP8 confocal microscope.

**Supplementary Table 2. Primary antibodies used in conventional immunofluorescence experiments.**

| Antibody | Dilution | Company | Catalogue number | Specie |
|---|---|---|---|---|
| Anti-$Na_V$1.5 | 1:50 | Alomone | ASC-005-GP | Guinea pig polyclonal |
| Anti-α-Actinin | 1:200 | Sigma | A7732 | Mouse monoclonal |
| Anti-SAP97 | 1:200 | Invitrogen | PA1-741 | Rabbit polyclonal |
| Anti-α1-Syntrophin | 1:200 | Invitrogen | PA5-22357 | Rabbit polyclonal |

**Supplementary Table 3. Secondary antibodies used in conventional immunofluorescence experiments.**

| Antibody | Dilution | Company | Catalogue number | Specie |
|---|---|---|---|---|
| Anti-Rabbit Alexa Fluor 488 | 1:500 | Invitrogen | A11034 | Goat anti-rabbit IgG |
| Anti-Mouse Alexa Fluor 568 | 1:500 | Invitrogen | A11031 | Goat anti-mouse IgG |
| Anti-*Guinea pig* DyLight 680 | 1:300 | Invitrogen | SA5-10098 | Goat anti-Guinea pig IgG |

**Cardiomyocytes plating and fixation.** Laminin-coated coverslips (rounded 13mm-diameter) were prepared by diluting the original stock of laminin (1mg/ml, Corning) 1:10 in PBS. After 1-2 hours of covering, we removed the excess diluted laminin on the top of the coverslips and the cardiomyocytes were left to adhere for 1-2 hours at RT. After that time, they were washed with PBS 1x and fixed using 4% PFA for 10min at RT. We washed the already fixed cardiomyocytes twice with PBS 1x and we followed the staining procedure for expansion microscopy as will be explained in the coming section.

**Expansion microscopy.** After the fixation, cells were permeabilized with 0.5% Triton-X100 in PBS for 15min at RT; followed by blocking in 5% BSA diluted in washing buffer (PBS+0.02% Tween 20) for 1 hour at RT. For expansion microscopy, we used the primary antibodies more concentrated than for conventional confocal microscopy (**Supplementary Table 4**); they were diluted in the blocking solution and incubated O/N at 4ºC. The next day, after three washes in the washing solution, the secondary antibodies (**Supplementary Table 5**) were diluted in blocking buffer and incubated for 2 hours at RT, followed again by two washes with washing solution and two more with

PBS. We used the crosslinker 10mg/ml 6-((Acryloyl)amino)hexanoic acid succinimidyl ester (or AcX) in PBS O/N at RT. The next step was the gelation procedure. We washed the coverslips twice with PBS and prepared in ice the following gelation solution: 4M Sodium Acrylate, 40% Acrylamide, 2% Bis-Acrylamide, PBS 10x, 10% TEMED and 10% APS. A drop of this solution was added in a gelation chamber per sample, we flipped each coverslip on them, and we left them in the incubator (37ºC) for 1 hour. After the gelation, we prepared a digestion buffer containing TAE buffer, 10% Triton-X100 in PBS, 5M NaCl, Proteinase K (26µl per every 2ml) and 1:400 DAPI. We moved the coverslips with the thin gel on their top from the gelation chambers to 12-multiwell plates with the digestion solution, and we let them digest for 4 hours at 37ºC. With the digestion the gels separated from the coverslips and increased their size three times. We moved them from the 12-multiwell plate to a 150-mm diameter culture plate per condition to add MilliQ water and let them expand O/N at RT. The next day they were imaged using a Leica Stellaris 8 microscope.

**Supplementary Table 4. Primary antibodies used in expansion microscopy experiments.**

| Antibody | Dilution | Company | Catalogue number | Specie |
|---|---|---|---|---|
| Anti-Na$_V$1.5 | 1:25 | Alomone | ASC-005-GP | *Guinea pig* polyclonal |

**Supplementary Table 5. Secondary antibodies used in expansion microscopy experiments.**

| Antibody | Dilution | Company | Catalogue number | Specie |
|---|---|---|---|---|
| Anti-*Guinea pig* Biotium CF647 | 1:100 | Biotium | 20108 | Goat anti-*Guinea pig* IgG |

**Western blot and membrane fractionation procedures.** To analyze total expression levels, we excised ventricles from untransfected, Kir2.1$^{WT}$, Kir2.1$^{D172N}$, Kir2.1$^{E299V}$, and Kir2.1$^{M301K}$ mice, and lysed them in ice-cold RIPA buffer (150mM NaCl, 10mM Tris-HCl, 1mM EDTA, 1% Triton X-100, 0.1% SDS and 0.1% Sodium deoxycholate). We sonicated the samples for 15s, 30% power, and conserved them in ice for 30min, applying 10s of vigorous shaking every 10min. We centrifuged the samples for 20min at 21000g and then recovered the supernatant. We quantified the resulting samples using Pierce™ BCA Protein Assay Kit (Thermo Fisher) and 25-80µg of protein resolved in 5-10% SDS-PAGE gels. We carried out the transference with the Trans-Blot Turbo Transfer System (Bio-Rad) (9min, 1.3A and 25V for mid-weight proteins; 13min, 1.3A and 25V for high-weight proteins). We blocked the membranes with 5% non-fat dry milk (diluted in PBS-0.05% Tween) when studying the $Na_V1.5$ expression, and with 5% BSA (diluted in PBS-0.05% Tween) for the remaining experiments, for 1 hour at RT. The employed antibodies and their concentrations are listed in **Supplementary Tables 6** and **7**.

To determine the Kir2.1 and/or $Na_V1.5$ protein levels at the plasma membrane, we used isolated mouse cardiomyocytes and followed the manufacturer's instructions for the Plasma Membrane Protein Extraction Kit (Abcam, ab65400). Briefly, we separated the cytoplasmic fraction and, after different centrifugations, the fraction containing plasma membrane proteins was pelleted and resuspended using the homogenization buffer. The enriched plasma membrane fraction was sonicated and kept in ice for 30min. The protein concentration in each fraction was quantified using Pierce™ BCA Protein Assay Kit (Thermo Fisher). We resolved total, cytosolic, and plasma membrane fractions of three animals per condition in 4-15% (gradient) Mini-PROTEAN TGX Stain-Free Gels (Bio-Rad). The transference, blocking, and blotting were performed as specified above. Note that there was slight contamination between fractions, and consequently, residual expression of membrane proteins in the cytosolic fraction, and *vice versa*.

**Supplementary Table 6. Primary antibodies used in western blotting experiments.**

| Antibody | Dilution | Company | Catalogue number | Specie |
|---|---|---|---|---|
| Anti-Kir2.1 | 1:200 | Abcam | Ab109750 | Rabbit monoclonal |
| Anti-$Na_V1.5$ | 1:500 | Sigma | S0819 | Rabbit polyclonal |
| Anti-GAPDH | 1:2500 | Abcam | Ab8245 | Mouse monoclonal |
| Anti-$Na^+/K^+$ ATPase | 1:4000 | Abcam | Ab7671 | Mouse monoclonal |

**Supplementary Table 7. Secondary antibodies used in western blotting experiments.**

| Antibody | Dilution | Company | Catalogue number | Specie |
|---|---|---|---|---|
| Anti-Rabbit Alexa Fluor 680 | 1:10000 | Invitrogen | A21076 | Goat anti-rabbit IgG |
| Anti-Mouse Alexa Fluor 680 | 1:10000 | Invitrogen | A21057 | Goat anti-mouse IgG |
| Anti-Rabbit HRP | 1:1000 | Invitrogen | 65-6120 | Goat anti-rabbit IgG |

**qRT-PCR.** Heart samples from untransfected, AAV9-Kir2.1$^{WT}$ and Kir2.1$^{M301K}$ mice were homogenized with the Trizol procedure for RNA extraction (ThermoFisher, 15596026). RNA concentration was measured using the ultraviolet (UV) spectrophotometry at 260nm (Nanodrop 2000, Thermo Scientific). cDNA was obtained using the High-Capacity cDNA Reverse transcription Kit (Applied Biosystems, 4368814), avoiding DNA contamination with DNase treatment of RNA samples before the qRT-PCR. The PCR protocol for obtaining the cDNA was: 10min at 25ºC – 120min at 37ºC – 5min at 85ºC. qPCR was performed using SYBR Green PCR Master Mix (Takara, RR420) and applying the following protocol: 1) 95ºC for 10min; 2) 95ºC for 15s; 3) 60ºC for 1min; 4) go to step 2 and repeat 39 more cycles; 5) melt curve from 65 to 95ºC with 0.5ºC increments for 5s. Primer sequences used were listed in **Supplementary Table 8**. Each reaction was run in triplicates. Gene expression values were normalized to the average expression of the housekeeping gene GAPDH.

**Supplementary Table 8. Primers used in RT-qPCR experiments.**

| Protein (Gene) | Forward primer (5'-3')<br>Melting Temperature | Reverse primer (5'-3')<br>Melting Temperature |
|---|---|---|
| $Na_V1.5$ (*SCN5A*) | TGAGTACACCTTCACCGCCA<br>66.7 ºC | TGAAAGTTCGAAGAGCCGACAA<br>68.0 ºC |
| Glyceraldehyde-3-phosphate dehydrogenase (*GAPDH*) | ACACTCACTCTTCTACCTTTG<br>55.1 °C | CAAATTCATTGTCGTACCAG<br>58.1 °C |

**Serological analysis of cardiac, hepatic and renal parameters.** Following a maxillofacial puncture performed at 9am, approximately 200µl of blood was collected into serum separator tubes from 43-week-old untreated, untransfected animals. This was done to compare cardiac, hepatic, and renal serum parameters with those of 43-week-old male mice treated with spermidine and expressing either Kir2.1$^{WT}$ or Kir2.1$^{M301K}$. To obtain serum, blood samples were left to stand in an upright position for 30 minutes to allow proper clot formation, and then centrifuged for 15 minutes at 3000rpm. The resulting serum was collected from the supernatant into clean Eppendorf tubes for subsequent serological analysis.

**Statistical analysis.** We used GraphPad Prism software version 7.0 and 8.0. We applied the statistical tests after a normal (Gaussian) distribution analysis (Shapiro-Wilk normality test and equality of variances with F test). With 2 groups and normal distribution, comparisons were made using unpaired 2-tailed Student's t-test. When the data do not follow a Gaussian distribution, we apply the nonparametric Mann-Whitney test. Unless otherwise stated, we used one- or two-way ANOVA followed by the post hoc Tukey test for comparison among more than two groups. If data did not follow a normal distribution, the nonparametric equivalent Kruskal-Wallis test was performed. To compare the trend of the sodium activation and inactivation curves across different membrane voltages between groups, an Ordinary one-way ANOVA was applied. To exclude outliers from some data sets we performed Grubbs' test, also known as the ESD (extreme studentized deviate) method, to determine whether a given value was a statistically significant outlier from the rest. For ECG recordings, we used 10-30 mice per condition and the analysis were carried out by two blinded researchers; on intracardiac stimulation procedures, around 10 animals of each genotype were used; and about 10-15 cells from 3 mice were studied in patch-clamp experiments. Unless other was specified, “N” means number of mice and “n” means the samples or cells analyzed in each case. Data are expressed as mean ± SEM, and differences are considered significant at $p<0.05$ (*$p<0.05$; **$p<0.01$; ***$p<0.001$; ****$p<0.0001$).

#### FIGURE LEGENDS

**Supplementary Figure 1. Generation of cardiac-specific SQTS3 mouse models**. **A**, **i**, AAV9 constructs containing the cardiac troponin-T promoter (cTnT), the *KCNJ2* (WT or mutant) gene, the Tomato reporter gene, an internal ribosome entry site (IRES) and the poly-adenosine (pA) sequence, all flanked by inverted terminal repeats (ITRs). **ii**, Representative fluorescence images of untransfected, Kir2.1$^{WT}$ and Kir2.1$^{M301K}$ hearts (we visualized more than 3 hearts per group). **iii**, Top, Haematoxylin-Eosin staining shows the normal structure of untransfected, and Kir2.1$^{WT}$ and Kir2.1$^{M301K}$ ventricles. Bottom, tdTomato immunostaining of untransfected, Kir2.1$^{WT}$ and Kir2.1$^{M301K}$ ventricles. Scale bar, 1mm. **iv**, Immunohistochemistry of tdTomato confirmed the infection in Kir2.1$^{WT}$ and Kir2.1$^{M301K}$ ventricles. Different scale bar as indicated. **B**, Left, Kir2.1 total protein levels in untransfected, Kir2.1$^{WT}$ and Kir2.1$^{M301K}$ hearts (N=6-11 animals per condition). Right, Kir2.1 membrane expression level in Kir2.1$^{WT}$ and Kir2.1$^{M301K}$ ventricular cardiomyocytes. T, C, M indicate total, cytosol, and membrane-enriched fraction, respectively (N=2 mice per condition). One-way ANOVA and unpaired 2-tailed Student's t-test were applied for statistics.

**Supplementary Figure 2. Echocardiographic analysis of Kir2.1$^{WT}$ and Kir2.1$^{M301K}$ mice reveals normal cardiac structure and function**. Values for Kir2.1$^{WT}$ (blue, N=10) and Kir2.1$^{M301K}$ (purple, N=10) are presented. **A**, Tricuspid Annular Plane Systolic Excursion (TAPSE), measured as distance in cm, analysis shows no right ventricular function defects. **B**, Interventricular septal end diastole (IVSd) is indicated. **C**, Ejection fraction and fractional shortening presented as percentages for all groups. Unpaired 2-tailed Student's t-test was applied to compare Kir2.1$^{M301K}$ mice *vs* Kir2.1$^{WT}$ animals.

**Supplementary Figure 3. AAV9-Kir2.1$^{WT}$ and untransfected mice behave similarly in terms of ECG, arrhythmia inducibility, and electrophysiology. A**, Upper panels, and from left to right, QT, QTc (Bazett's correction), and QTc (Framingham's formula) in Kir2.1$^{WT}$ (blue; N=10-28) *vs* untransfected (black; N=10-29) mice. Lower panels, and from left to right, PR interval, QRS complex, and P wave duration in Kir2.1$^{WT}$ and untransfected mice. Mann-Whitney test (QT/QTc intervals, PR intervals and QRS complexes) and unpaired 2-tailed Students't-test (P waves) were used. **B**, Left, ECG recordings during PES protocols. Scale bar, 500ms. Right, number of Kir2.1$^{WT}$ and untransfected mice with >500ms atrial and ventricular arrhythmias. Fisher's exact test was applied. **C-E**, Adapted from Macías, A *et al.*[11] The exact sample size for each experiment is indicated in the corresponding panel. **C**, IV relationships of $I_{K1}$, and **D**, $I_{Na}$ IV recorded from non-infected (black), Kir2.1WT (blue), and Kir2.1$^{\Delta 314-315}$ (red). Two-way

ANOVA, followed by Tukey's multiple comparisons, were applied. **E**, Representative AP recordings show no differences between Kir2.1$^{WT}$ and non-infected cardiomyocytes in terms of APD and RMP. Different colors in the same group identify cells coming from one animal. Note that "non-infected mice" refers to "untransfected animals". Statistical analyses were conducted using two-level hierarchical one-way ANOVA analysis, followed by Bonferroni's post-test.

**Supplementary Figure 4. Ventricular refractory periods are shorter in Kir2.1$^{M301K}$ than Kir2.1$^{WT}$ mice**. Ventricular refractory period of Kir2.1$^{WT}$ (N=8, blue), and Kir2.1$^{M301K}$ (N=8, purple) (65.5±1.9ms and 40.1±3.5ms, respectively). Unpaired 2-tailed Students' t-test was used for statistical comparisons (****p<0.0001).

**Supplementary Figure 5. Kir2.1$^{M301K}$ reduces ventricular APD at all stimulation frequencies**. For ventricular cardiomyocytes isolated from Kir2.1$^{WT}$ (blue; N=3, n=9-16) and Kir2.1$^{M301K}$ (purple; N=3, n=14-18) hearts, APDs at 20 ($APD_{20}$), 50 ($APD_{50}$), and 70 ($APD_{70}$) % of repolarization at 1, 2, 4, 5 and 10 Hz are shown on different and consecutive panels (**A-C**, respectively). Mann-Whitney tests were applied (****p<0.0001, ***p<0.001, and *p<0.05).

**Supplementary Figure 6. Examples of the subtraction process to obtain barium ($BaCl_2$)-sensitive $I_{K1}$ current.** Representative traces of total $K^+$ currents (top), $K^+$ current in presence of 500µM $BaCl_2$ (middle), and $BaCl_2$-sensitive $K^+$ currents (bottom) from ventricular Kir2.1$^{WT}$ (blue) and Kir2.1$^{M301K}$ (purple) cardiomyocytes. Currents were elicited by 500ms voltage steps (each 10mV) ranging from -140 to 0 mV from a holding potential of -80mV. Time (ms) is on the X axis, and current, in pA, on the Y axis. Dashed lines highlight the baseline (0pA).

**Supplementary Figure 7. The mutation Kir2.1$^{M301K}$ slightly changes towards hyperpolarized voltages the $I_{Na}$ activation properties**. **A**, Graphs show sodium activation (<u>left</u>) and inactivation (<u>right</u>) parameters ($V_{50}$ and slope) for Kir2.1$^{WT}$ (blue; N=3, n=14-20) and Kir2.1$^{M301K}$ (N=3; n=13-16). Mann-Whitney test was applied for comparisons.

**Supplementary Figure 8. Main Kir2.1-$Na_V$1.5 interactors distribution pattern is not modified in Kir2.1$^{M301K}$ ventricular cardiomyocytes.** Representative confocal images of SAP97 (purple), α1-Syntrophin (yellow) and α-Actinin (light blue) staining in Kir2.1$^{WT}$ (**A**) and Kir2.1$^{M301K}$ (**B**) ventricular cardiomyocytes showing the normal distribution pattern of these proteins. Scale bars, 5µm.

**Supplementary Figure 9. Weight loss quantification while administering 10mg/Kg spermine per day to Kir2.1$^{WT}$ and Kir2.1$^{M301K}$ animals.** Mice's weight is expressed as percentage to show the overall decrease over time. The animals started the treatment with 30.7±1.2g of weight. Sixteen animals per condition (Kir2.1$^{WT}$ in blue, and Kir2.1$^{M301K}$ in purple) were under the treatment until day 10, and 8 mice per genotype until day 20.

**Supplementary Figure 10. Spermine administration prolonged APD in Kir2.1$^{M301K}$ ventricular cardiomyocytes. A**, from left to right, PR interval, QRS complex and RR interval durations in Kir2.1$^{WT}$ (blue; N=8-16) and Kir2.1$^{M301K}$ (purple; N=8-16) over the time after starting the treatment with spermine. **B**, APD at different repolarization percentages (20, 50, 70 and 90) at 1Hz (left) and 5Hz (right), and **C**, RMP (mV), recorded from WT control (dark blue; N=3; n=9-11), M301K control (dark purple; N=3; n=8-9), M301K 10-day spermine (light purple; N=3; n=10-14), and M301K 20-day spermine (dark pink; N=3, n=11) ventricular cardiomyocytes. Asterisks in the graphs in panel B indicate statistically differences between each experimental condition (differentiated by color) and the M301K control group. **D**, Activation and inactivation curves of $I_{Na}$, and **E**, activation (left) and inactivation (right) parameters ($V_{50}$ and slope), recorded from WT control ( N=3; n=14-18), M301K control (N=3; n=13-16), M301K 10-day spermine (N=3; n=8-11), and M301K 20-day spermine (N=3, n=8-10) cells. The exact sample size for each experiment is indicated in the corresponding panels and spermine is abbreviated as "SPM". Two-way ANOVA, Mann-Whitney tests and Ordinary one-way ANOVA were applied for comparisons (****p<0.0001, **p<0.01, and *p<0.05).

**Supplementary Figure 11. The treatment with spermine did not modify significantly neither the AP characteristics, $I_{K1}$ nor $I_{Na}$ in Kir2.1$^{WT}$ ventricular cardiomyocytes. A**, Representative traces of APs recorded from WT control (blue), WT 10-day spermine (sea blue), and WT 20-day spermine (green) cardiomyocytes paced at 1Hz. **B**, AP parameters: RMP (mV), *d*V/*d*t (V/s), and AP amplitude (mV) recorded from WT control (N=3; n=11), WT 10-day spermine (N=3; n=13), and WT 20-day spermine (N=3; n=11) myocytes. **C**, APD at different repolarization percentages (20, 50, 70 and 90) at 1Hz (left) and 5 Hz (right) recorded from WT control (N=3; n=8-10), WT 10-day spermine (N=3; n=11), and WT 20-day spermine (N=3; n=8-10) ventricular cardiomyocytes. Slight green asterisk indicates the statistical difference between WT 20-day spermine and WT control. **D**, $APD_{90}$ at 1 (left) and 5 Hz (right) are shown for each group following the same color pattern. **E-H**, Outward $I_{K1}$ (**E**), $I_{Na}$ IVs (**F**), and activation and inactivation curves (**G**) and parameters ($V_{50}$ and slope) (**H**) of $I_{Na}$ recorded from WT control, WT 10-day spermine, and WT 20-day spermine cardiomyocytes. For **E-H**, color pattern and sample size are indicated in the corresponding panels, and spermine is abbreviated as "SPM". Two-way

ANOVA, Mann-Whitney tests and Ordinary one-way ANOVA were applied for comparisons (*p<0.05).

**Supplementary Figure 12. Chronic administration of spermidine in Kir2.1$^{M301K}$ mice prolonged their APD to achieve WT levels. A**, from left to right, PR interval, QRS complex, and RR interval durations in Kir2.1$^{WT}$ (blue) and Kir2.1$^{M301K}$ (purple) over the time after starting the treatment with spermidine. The sample size is indicated in every time point. **B**, APD at different repolarization percentages (20, 50, 70 and 90) at 1Hz (left) and 5Hz (right), and **C**, RMP (mV), recorded from WT control (dark blue; N=3; n=8-10), M301K control (dark purple; N=3; n=8-9), M301K 20-day spermidine (dark pink; N=3; n=9), and M301K 6-month spermidine (light pink; N=3, n=8-11) ventricular cardiomyocytes. Asterisks in the graphs in panel B indicate statistically differences between each experimental condition (differentiated by color) and the M301K control group. **D**, Activation and inactivation curves of $I_{Na}$, and **E**, activation (left) and inactivation (right) parameters ($V_{50}$ and slope), recorded from WT control (N=3; n=14-18), M301K control (N=3; n=13-16), M301K 20-day spermidine (N=3; n=8-11), and M301K 6-month spermidine (N=3, n=8-10) cells. Spermidine is indicated as "SPD" through the panels. Two-way ANOVA, Mann-Whitney tests, and Ordinary one-way ANOVA were applied for comparisons (****p<0.0001, ***p<0.001, **p<0.01, and *p<0.05).

**Supplementary Figure 13. Chronic administration of spermidine prolonged the APD in Kir2.1$^{WT}$ myocytes due to a reduction in their $I_{K1}$ outward current. A**, Representative traces of APs recorded from WT control (blue), WT 20-day spermidine (light green), and WT 6-month spermidine (dark green) cardiomyocytes paced at 1Hz. **B**, AP parameters: RMP (mV), *d*V/*d*t (V/s), and AP amplitude (mV) recorded from WT control (N=3; n=11), WT 20-day (N=3; n=6), and WT 6-month spermidine (N=3; n=6-8) myocytes. **C**, APD at different repolarization percentages (20, 50, 70 and 90) at 1Hz (left) and 5Hz (right) recorded from WT control (N=3; n=8-10), WT 20-day spermidine (N=3; n=5), and WT 6-month spermidine (N=3; n=6-7) ventricular cardiomyocytes. Asterisks indicate statistically differences between each experimental condition (differentiated by color) and the WT control group. **D**, $APD_{90}$ at 1 (left) and 5 Hz (right) are shown for each group following the same color pattern. **E-H**, Outward $I_{K1}$ (**E**), $I_{Na}$ IVs (**F**), and activation and inactivation curves (**G**) and parameters ($V_{50}$ and slope) (**H**) of $I_{Na}$ recorded from WT control, WT 20-day spermidine, and WT 6-month spermidine myocytes. For **E-H**, color pattern and sample size are indicated in the corresponding panels, and spermidine is abbreviated as "SPD". Two-way ANOVA, Mann-Whitney tests and Ordinary one-way ANOVA were applied for comparisons (***p<0.001, **p<0.01, and *p<0.05).

**Supplementary Figure 14. Chronic treatment with spermidine in aged Kir2.1$^{M301K}$ mice also prolonged the QT/QTc intervals. A-B**, Left, QT interval duration (raw data in **A**) and corrected values by the Bazett's formula (**B**) in 81-week-old WT control (dark blue; N=2), WT treated for 6 month with spermidine (purple; N=6), M301K control (dark green; N=5), and M301K treated for 6 month with spermidine (light pink; N=5). Right, QT interval duration recorded from Kir2.1$^{WT}$ (blue; N=5) and Kir2.1$^{M301K}$ (purple; N=5) mice over time following the administration of 3mM spermidine via drinking water. The *p* values indicate the statistical significance between the linear regression of both groups in each case. **C**, from left to right, PR interval, QRS complex and RR interval durations in Kir2.1$^{WT}$ (blue; N=5) and Kir2.1$^{M301K}$ (purple; N=5) over time after starting the treatment with spermidine. In each graph, dashed lines indicate the linear regression of the ECG values analyzed over time (WT in light blue, and M301K in light purple). Two-way ANOVA and a linear regression analysis were applied for comparisons (****p<0.0001 and *p<0.05).

**Supplementary Figure 15. Long-term treatment with spermidine reduces significantly mice's body weight but does not result toxic on an organic level. A-B**, Body weight of 42-week-old (**A**) and 81-week-old (**B**) untreated (black; N=5-8), Kir2.1$^{WT}$ treated (dark green; N=5-6), and Kir2.1$^{M301K}$ treated (light pink; N=5) for 6 months with 3mM spermidine. **C**, Heart weight/body weight (HW/BW) ratio of 42-week-old animals both control (N=5) and treated with spermidine for 6 months (N=5). **D**, Haematoxylin-Eosin (H&E) and Masson's trichrome staining (Masson's) for liver, kidney, lung, and spleen tissue slices obtained from 42-week-old untransfected untreated (control), and Kir2.1$^{WT}$ and Kir2.1$^{M301K}$ 6 months treated mice. Spermidine is indicated as "SPD" along the panels. Mann-Whitney tests were applied for comparisons (**p<0.01 and *p<0.05).

**Supplementary Figure 16. Serological analysis of cardiac, hepatic and renal parameters reveals no abnormalities after 6 months of treatment with 3mM spermidine.** Biochemical parameters obtained from serum samples of untransfected untreated (control, in black; N=4), and Kir2.1$^{WT}$ (blue; N=6) and Kir2.1$^{M301K}$ (purple; N=5) treated for 6 months with spermidine, organized by profiles: **A**, cardiac profile, including C-reactive protein and creatinine; **B**, ions (calcium and phosphorus); **C**, hepatic profile, containing AST/GOT and ALT/GPT; **D**, renal profile, including total proteins, alkaline phosphatase, albumin, glucose, and ureic nitrogen. The serum was derived from blood samples obtained through maxillofacial puncture. Spermidine is indicated as "SPD" along the panels. Mann-Whitney and Kruskal-Wallis tests were applied for comparisons (*p<0.05).

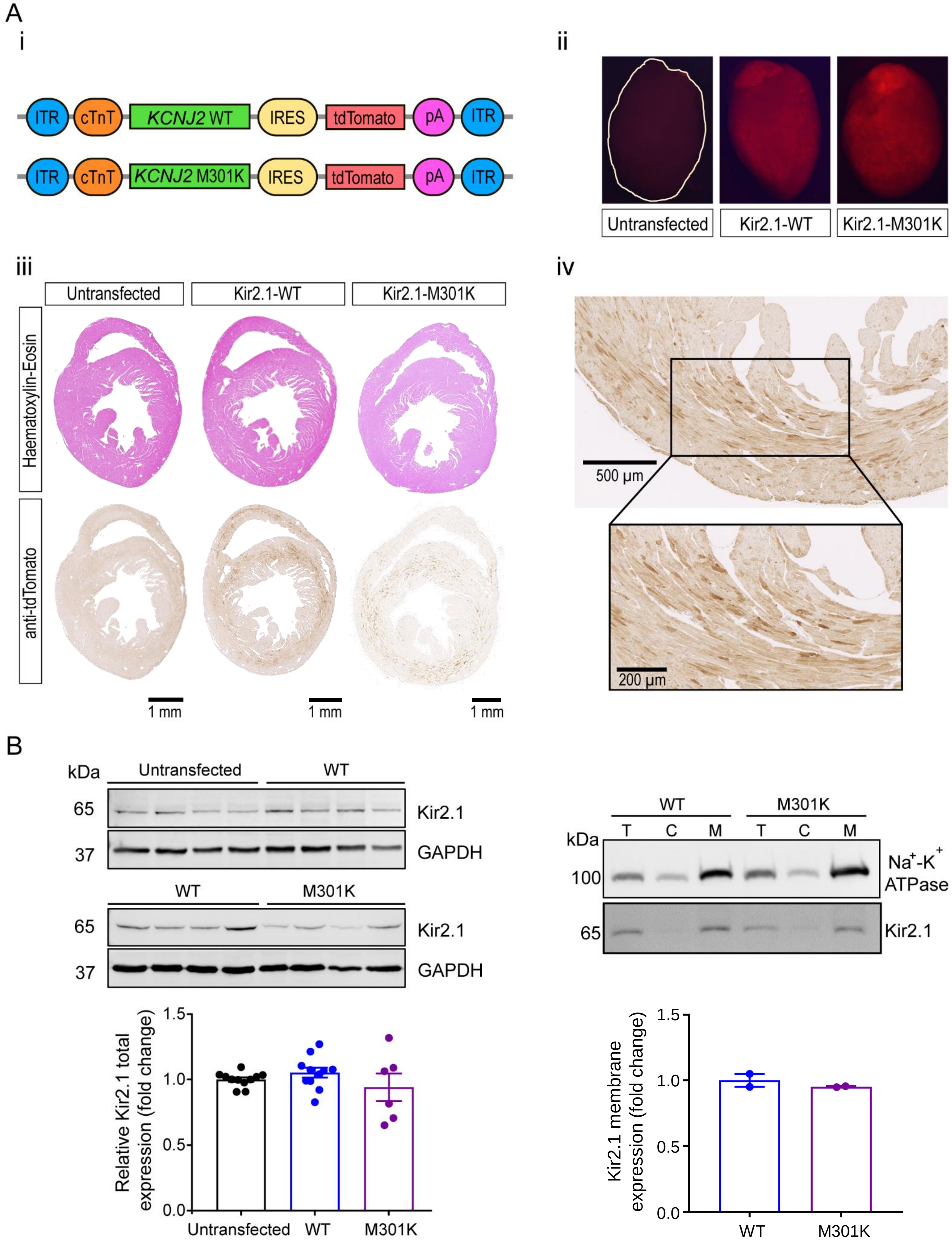


Supplementary Figure 1

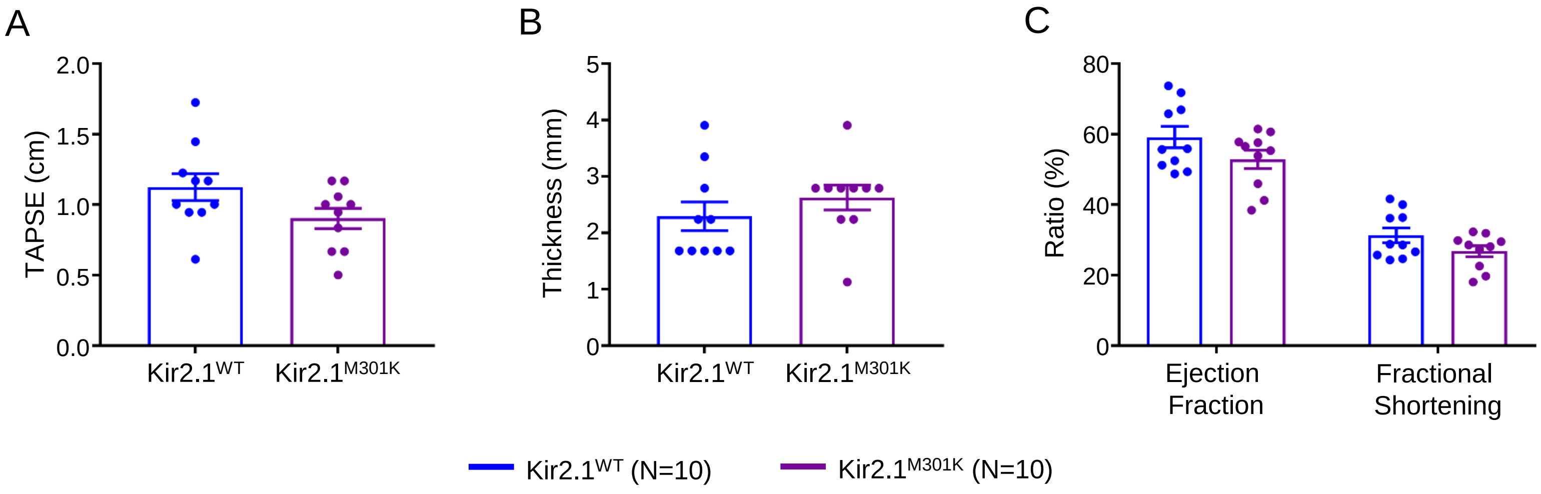


Supplementary Figure 2

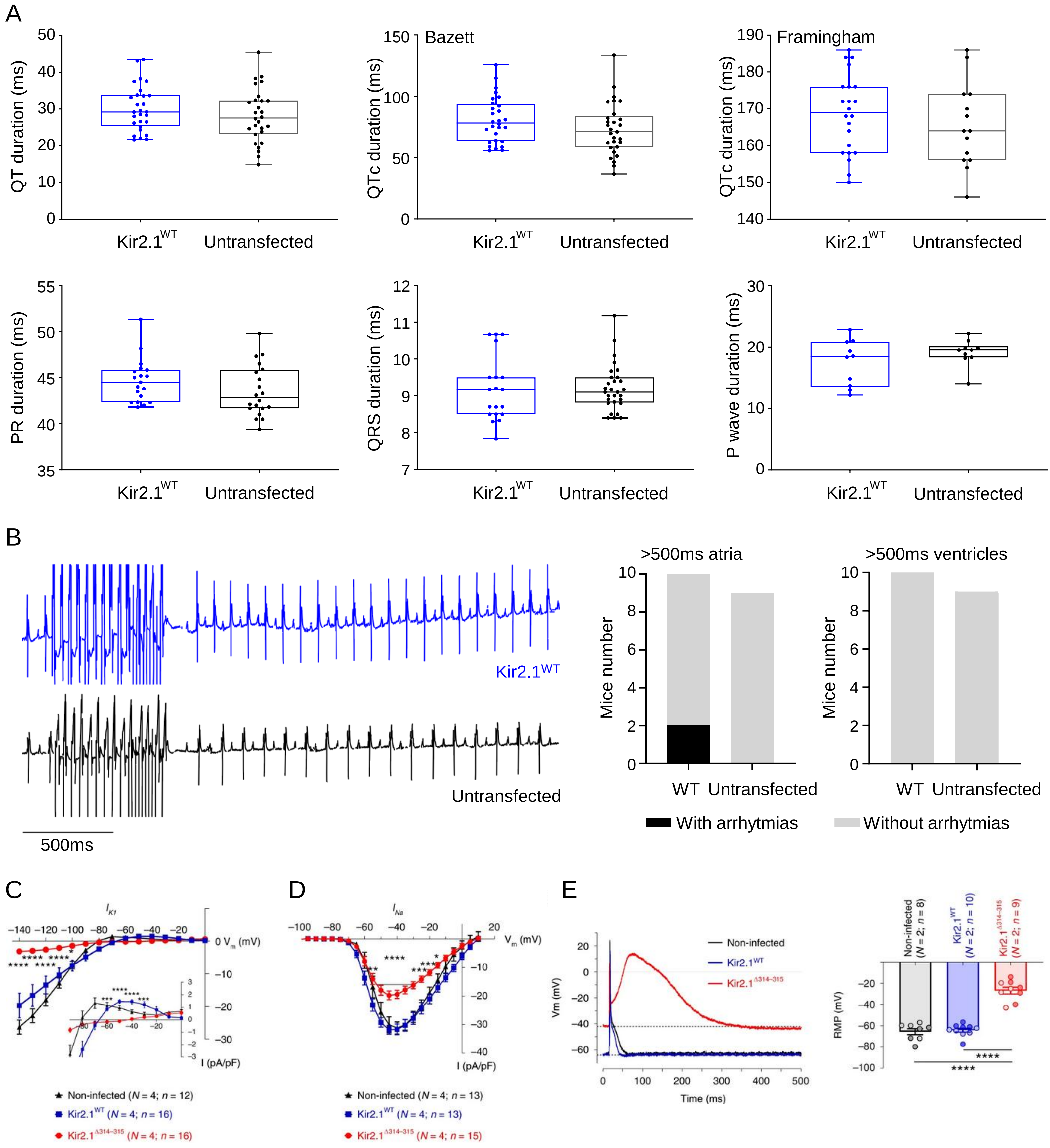


Supplementary Figure 3

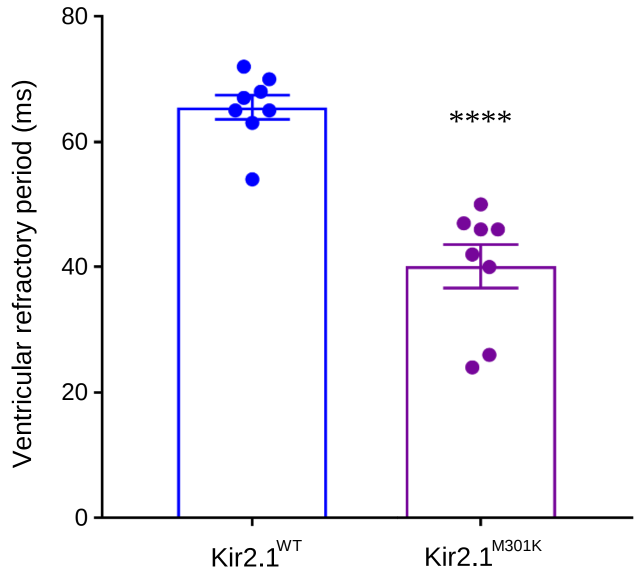


Supplementary Figure 4

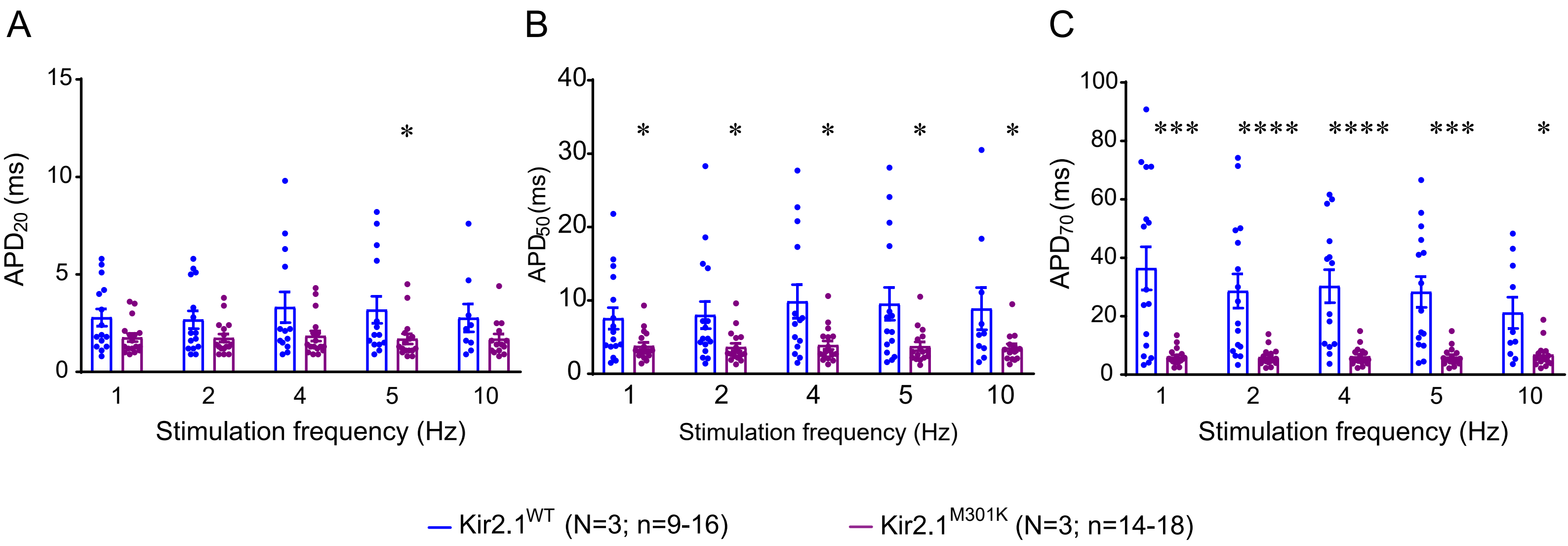


Supplementary Figure 5

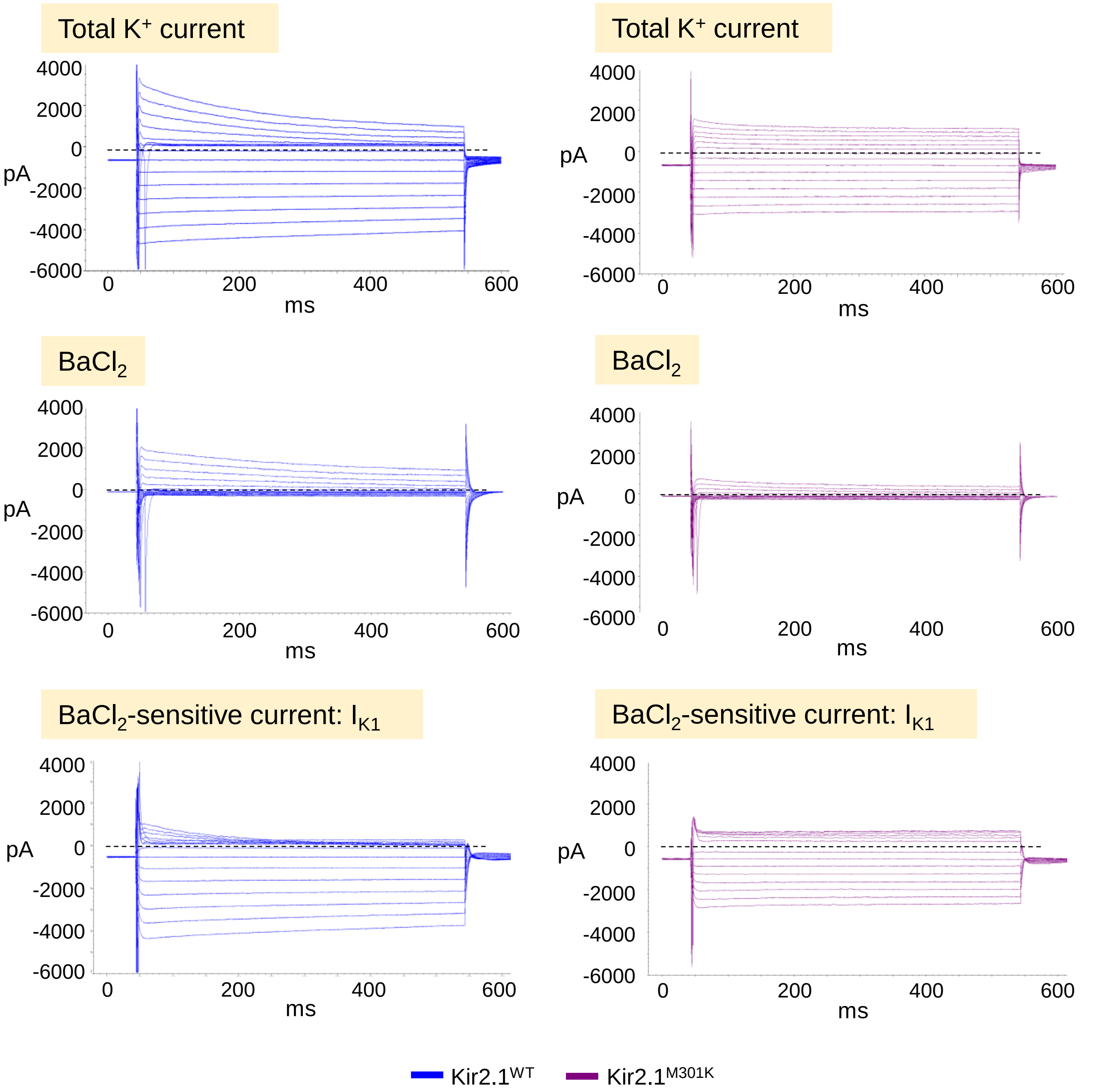


Supplementary Figure 6

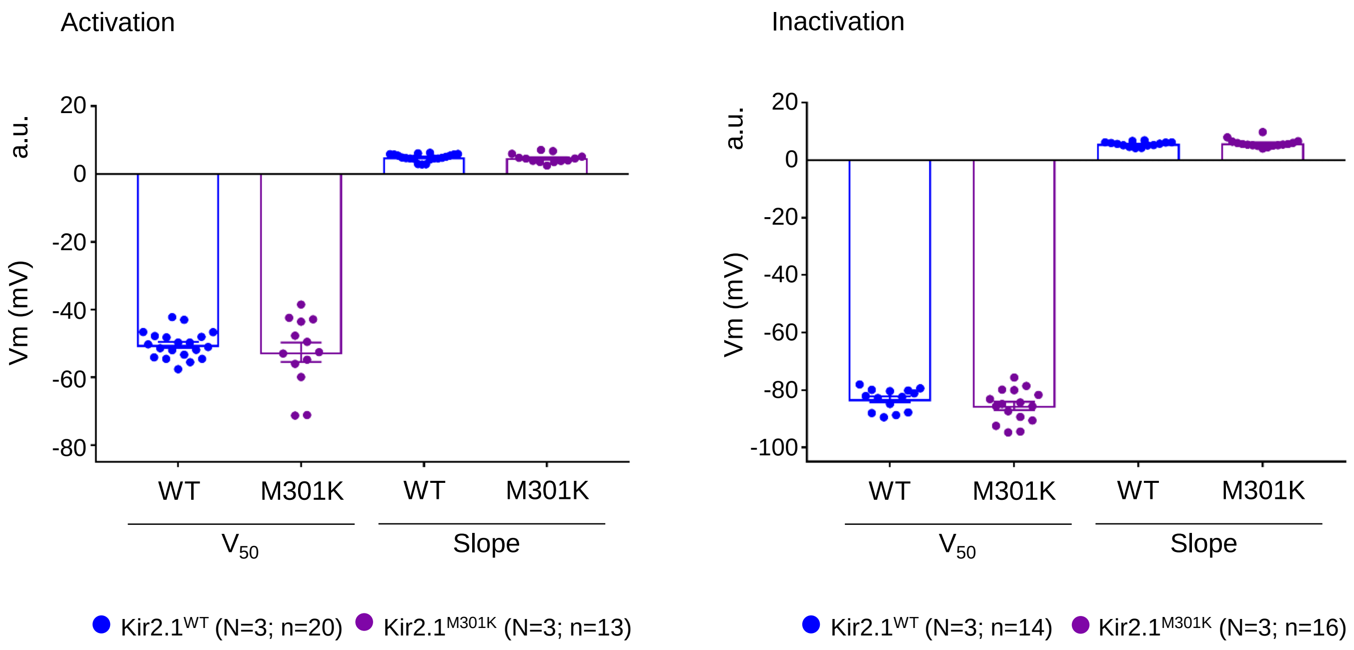


Supplementary Figure 7

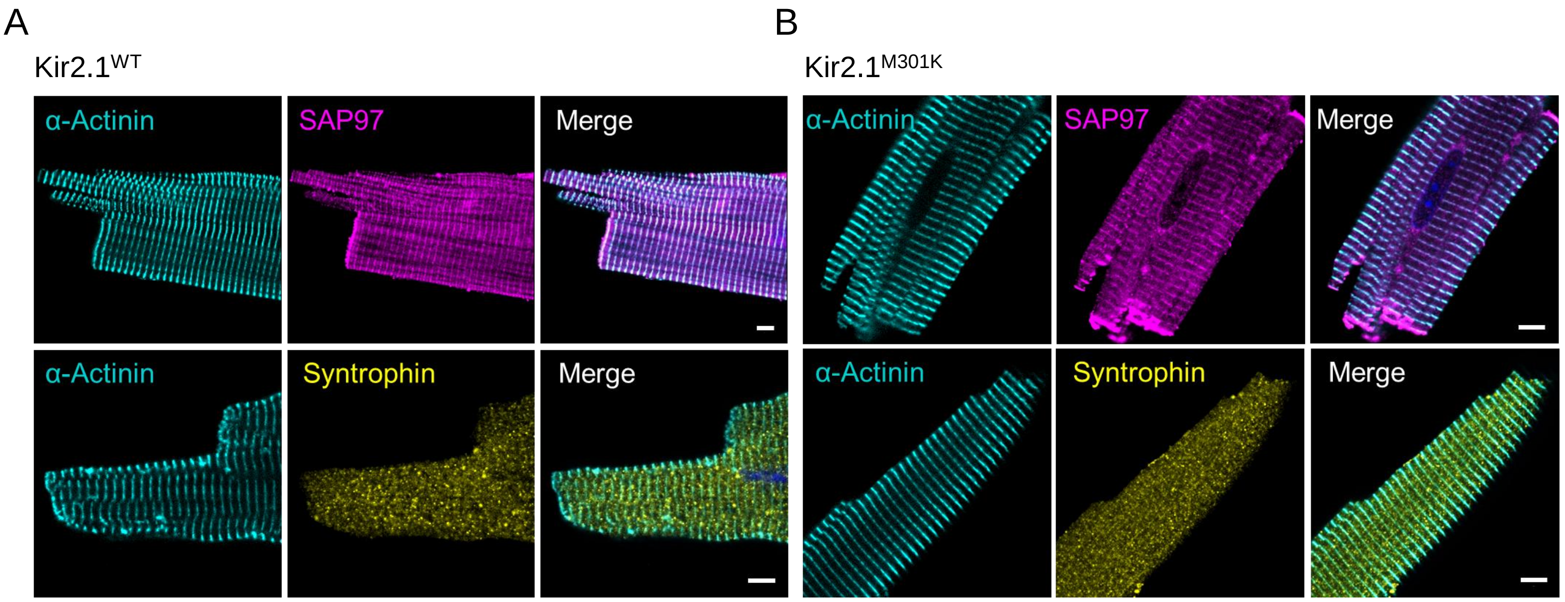


Supplementary Figure 8

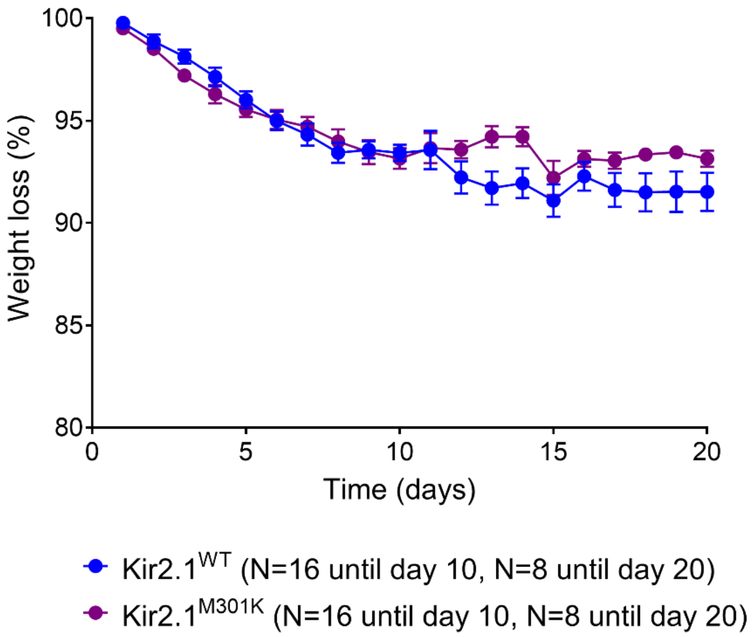


Supplementary Figure 9

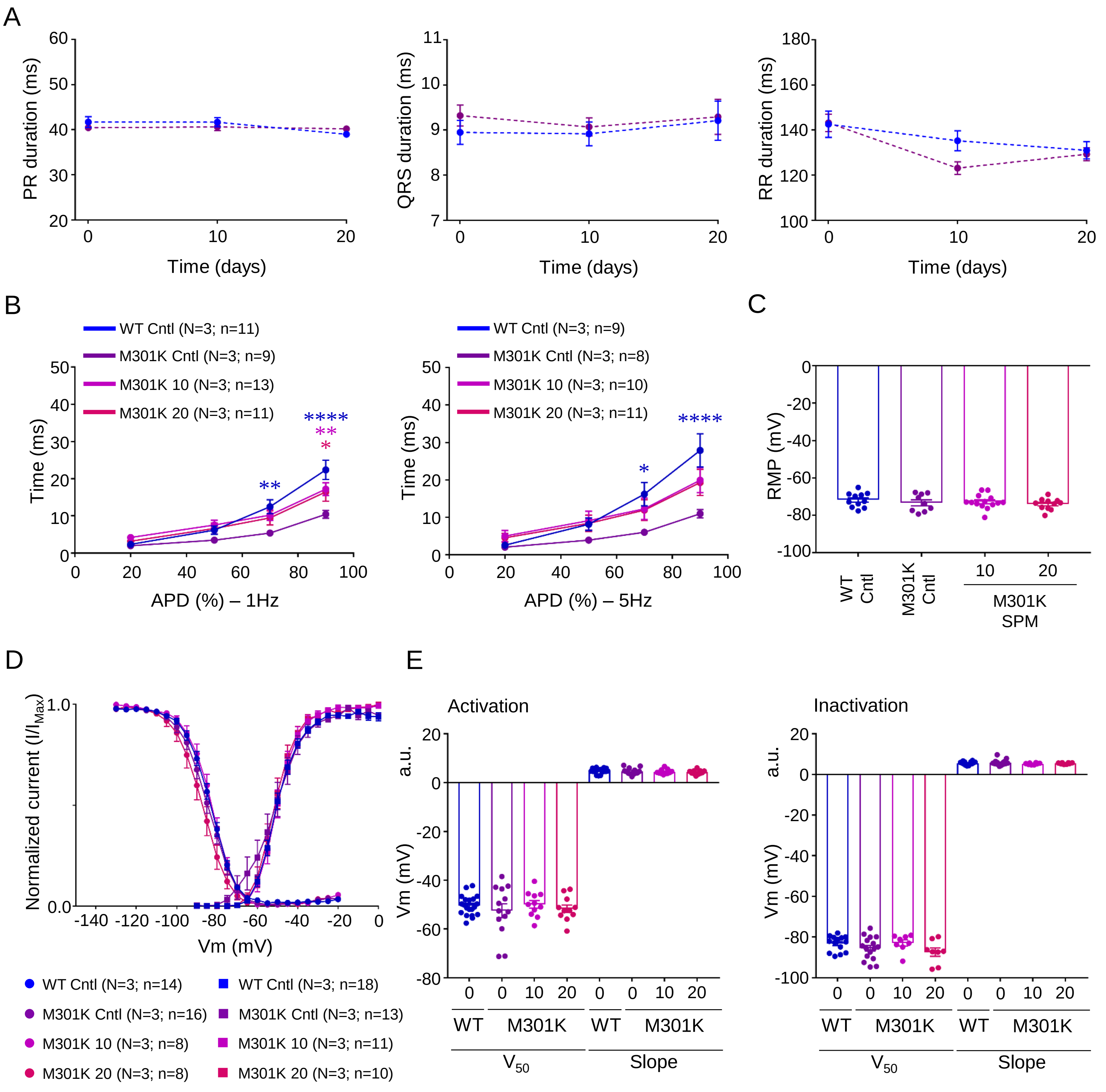


Supplementary Figure 10

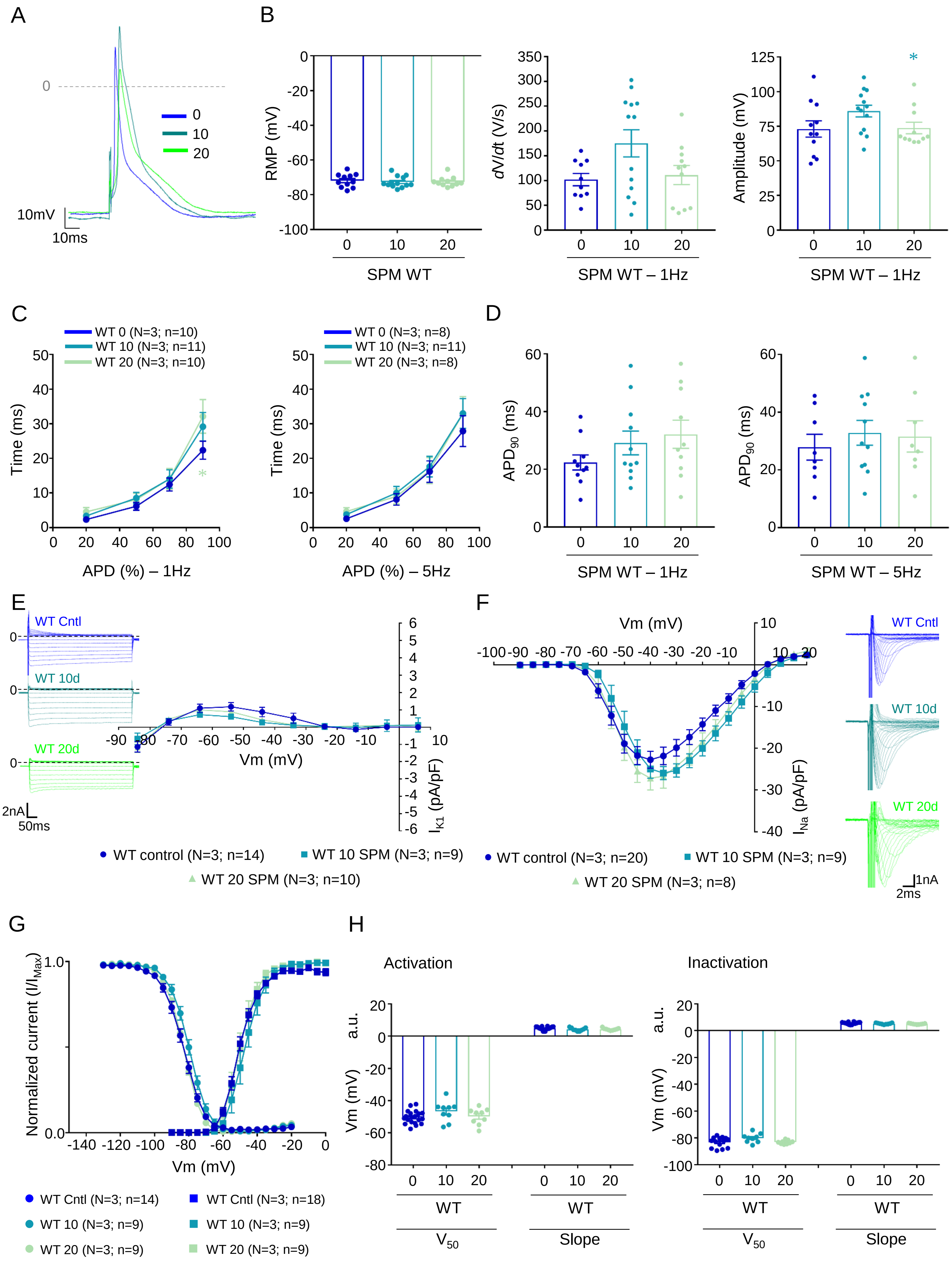


### Supplementary Figure 11

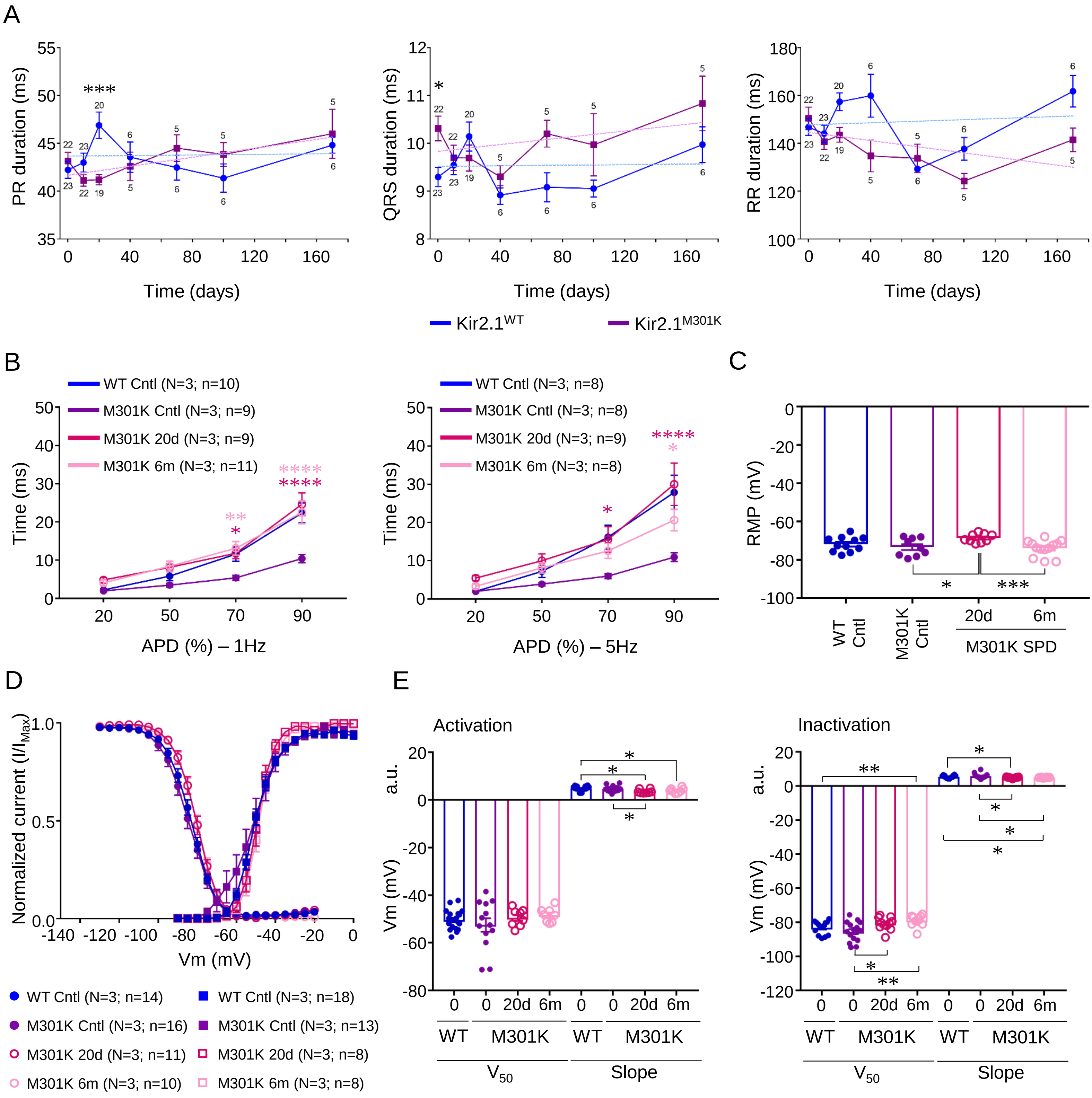


Supplementary Figure 12

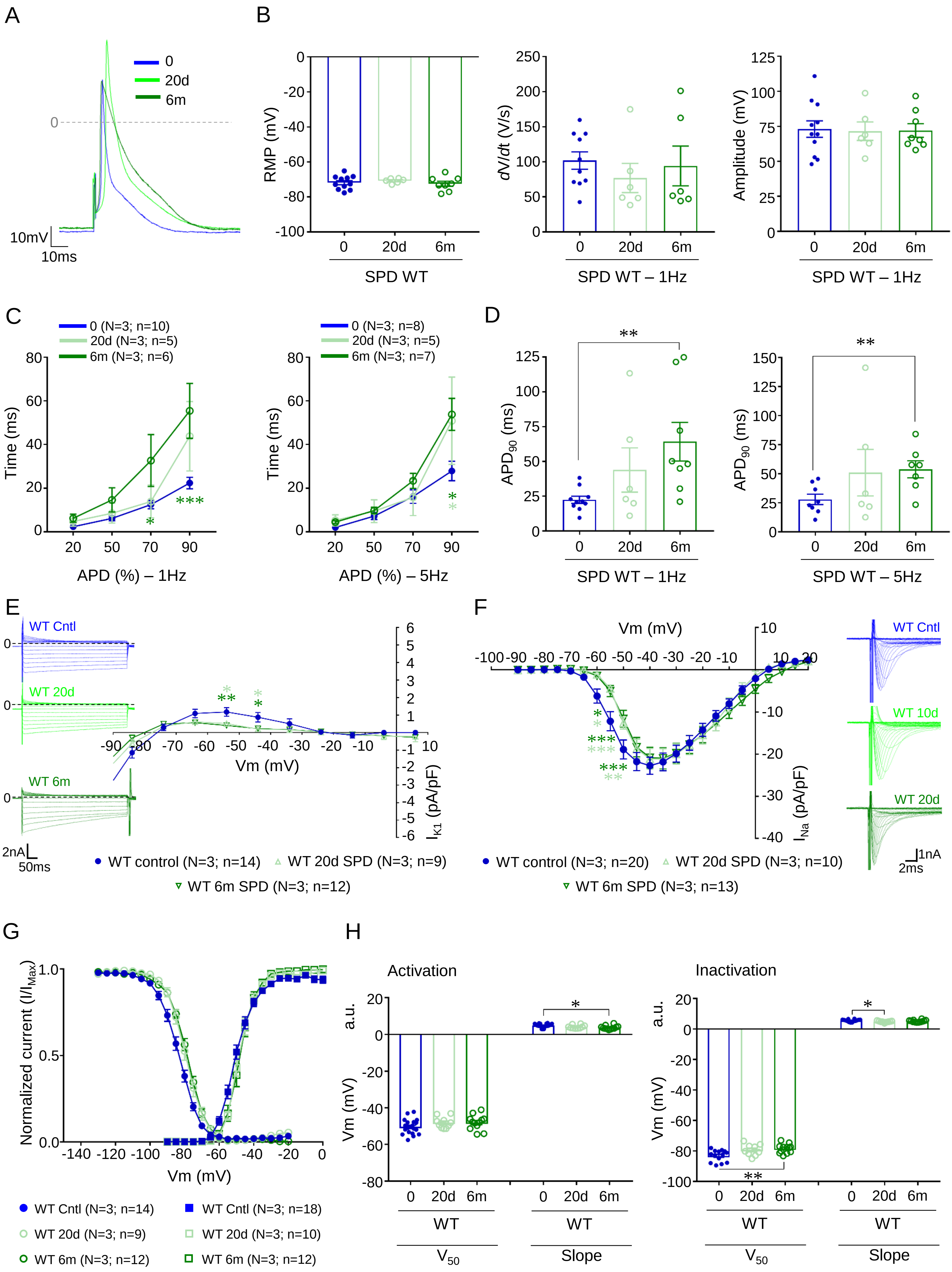


Supplementary Figure 13

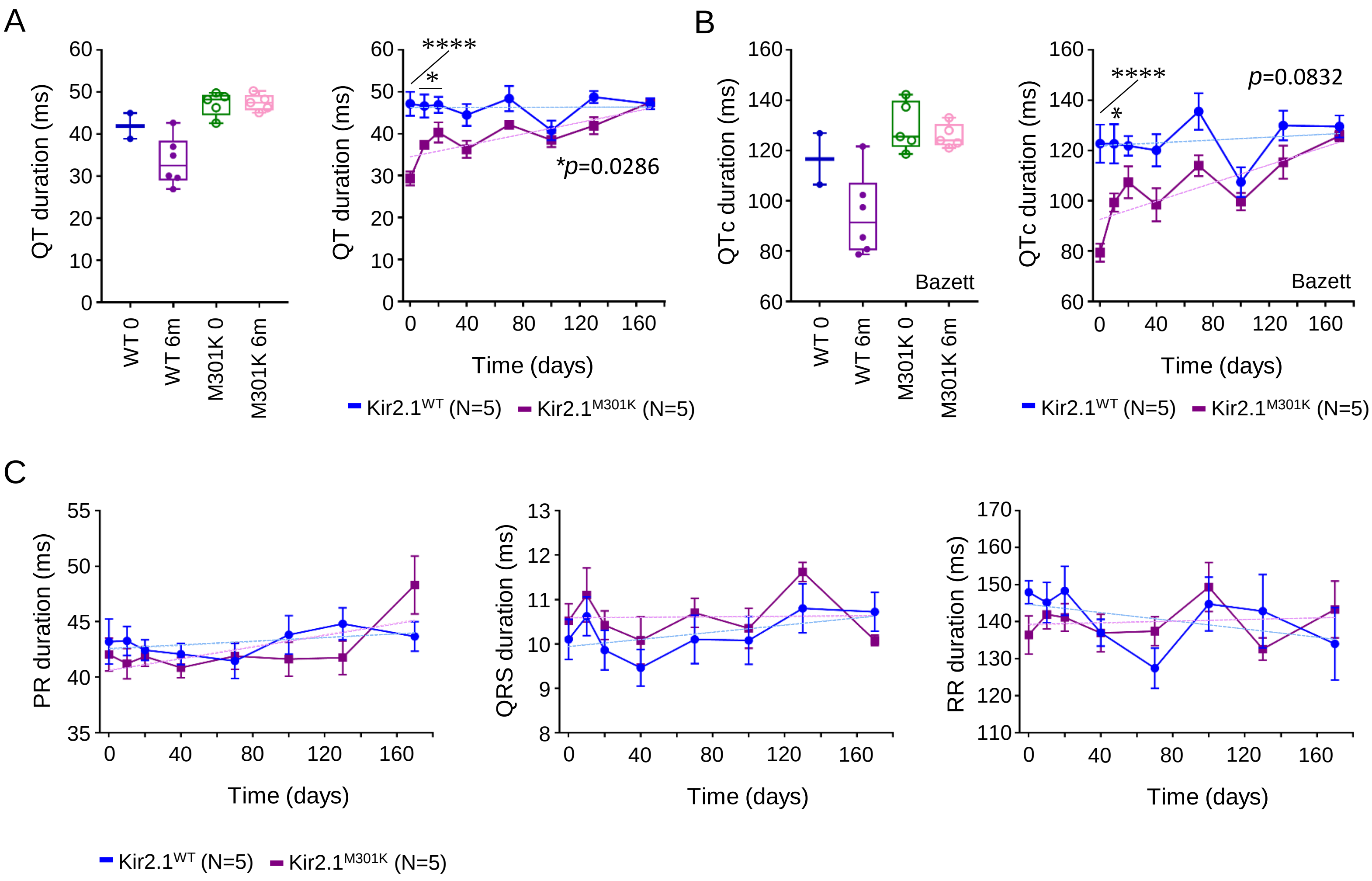


Supplementary Figure 14

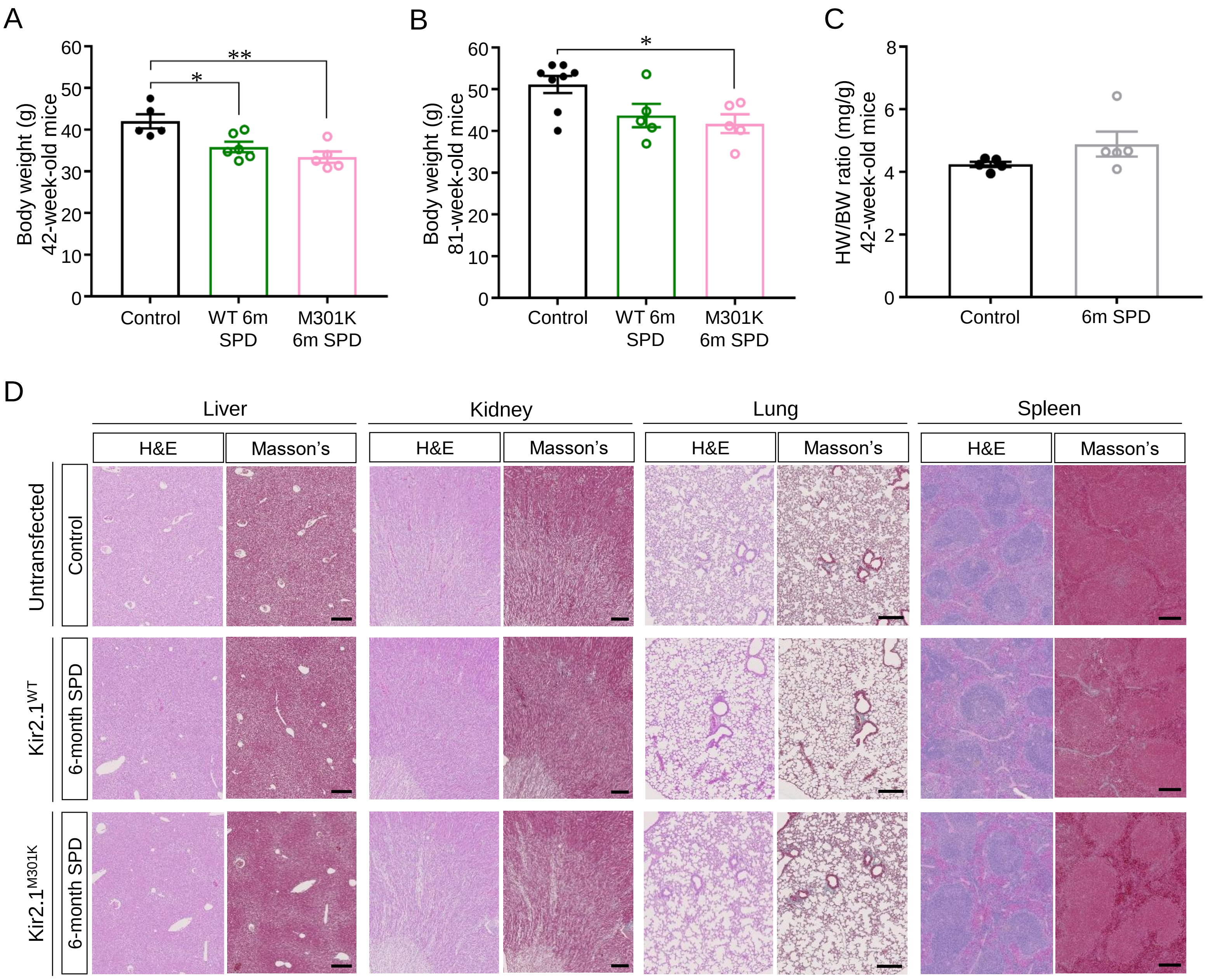


Supplementary Figure 15

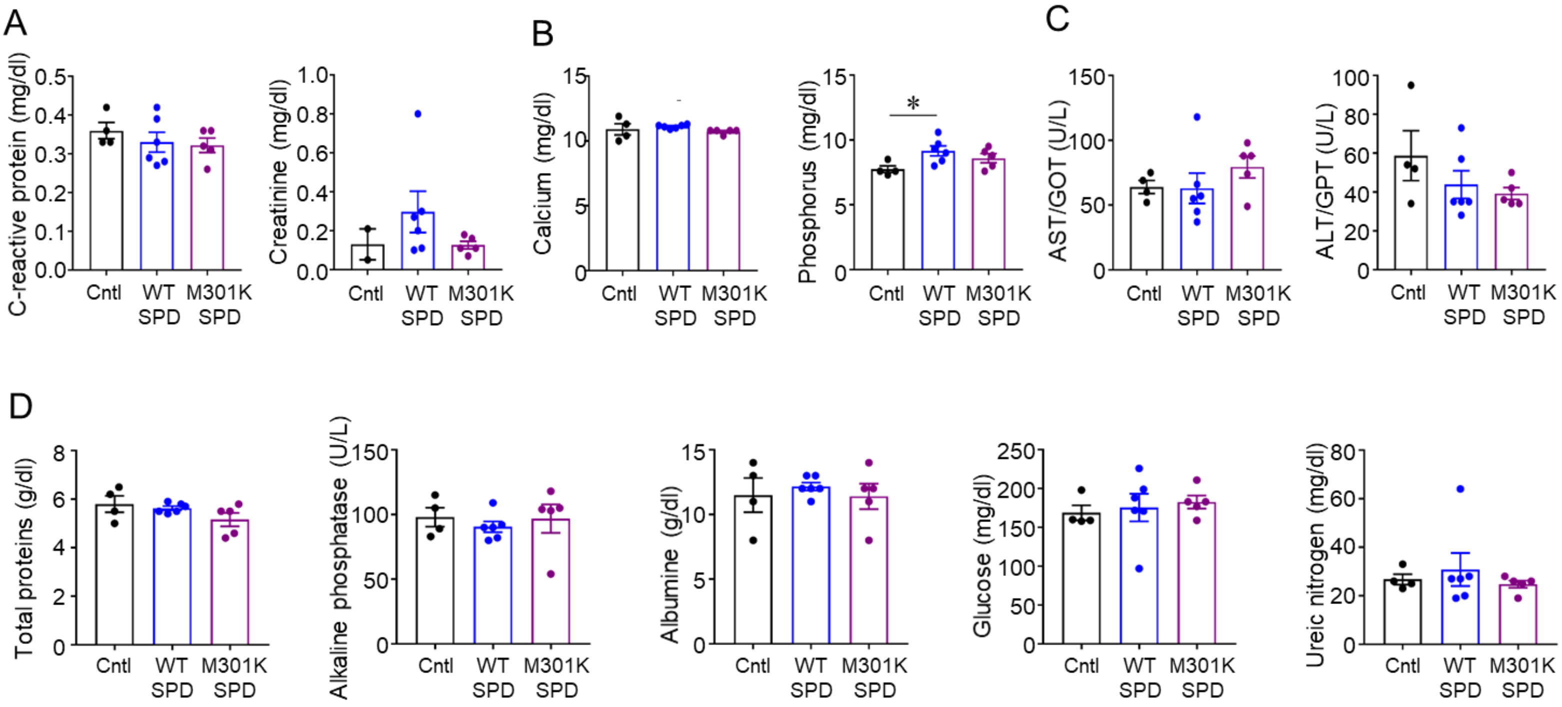


Supplementary Figure 16
